# The Zinc-BED transcription factor Bedwarfed promotes proportional dendritic growth and branching through transcriptional and translational regulation in *Drosophila*

**DOI:** 10.1101/2023.02.15.528686

**Authors:** Shatabdi Bhattacharjee, Eswar Prasad R. Iyer, Srividya Chandramouli Iyer, Sumit Nanda, Myurajan Rubaharan, Giorgio A. Ascoli, Daniel N. Cox

## Abstract

Dendrites are the primary points of sensory or synaptic inputs to a neuron and play an essential role in synaptic integration and neural function. Despite the functional importance of dendrites, relatively less is known about the underlying mechanisms regulating cell-type specific dendritic patterning. Herein, we have dissected functional roles of a previously uncharacterized gene, *CG3995*, in cell-type specific dendritic development in *Drosophila melanogaster*. *CG3995*, which we have named *bedwarfed* (*bdwf*), encodes a zinc-finger BED-type protein which is required for proportional growth and branching of dendritic arbors, exhibits nucleocytoplasmic expression, and functions in both transcriptional and translational cellular pathways. At the transcriptional level, we demonstrate a reciprocal regulatory relationship between Bdwf and the homeodomain transcription factor (TF) Cut. We show that Cut positively regulates Bdwf expression and that Bdwf acts as a downstream effector of Cut-mediated dendritic development, whereas overexpression of Bdwf negatively regulates Cut expression in multidendritic sensory neurons. Proteomic analyses revealed that Bdwf interacts with ribosomal proteins and disruption of these proteins produced phenotypically similar dendritic hypotrophy defects as observed in *bdwf* mutant neurons. We further demonstrate that Bdwf and its ribosomal protein interactors are required for normal microtubule and F-actin cytoskeletal architecture. Finally, our findings reveal that Bdwf is required to promote protein translation and ribosome trafficking along the dendritic arbor. Taken together, these results provide new insights into the complex, combinatorial and multi-functional roles of transcription factors (TFs) in directing diversification of cell-type specific dendritic development.

## Introduction

Neurons are highly complex, polarized cells that come in an astonishing number of shapes and sizes, attributable largely to their elaborate, cell-type specific dendritic arborization patterns that are adjusted to cover their receptive fields (Lefebvre, Sanes and Kay, 2015; Prigge and Kay, 2018). Since dendrites are primarily specialized to receive/process neuronal inputs, the specific morphology of the dendrite can govern neuronal function, signal integration, and circuit assembly. Thus, understanding the biological mechanisms regulating the growth and development of dendritic arbors is of particular significance for nervous system organization and function. Numerous studies have demonstrated that the acquisition of class-specific dendritic architectures is subject to regulation by complex genetic and molecular programs involving intrinsic factors and extrinsic cues including cytoskeletal regulation, transcriptional regulation, cell-signaling and cell-cell interactions (Tavosanis, 2021; Lefebvre, 2021).

Transcription factors (TFs) are one class of molecules that have emerged as critical regulators of dendritic development, where distinct or combinatorial TF activity has been shown to drive dendrite morphogenesis and cell-type specific dendritic diversity (Jan and Jan, 2010; de la Torre-Ubieta and Bonni, 2011; Santiago and Bashaw, 2014; Valnegri, Puram and Bonni, 2015; Nanda et al., 2017; Pai and Moore, 2021). In the mammalian brain, studies have also shown that the morphological identity of neurons in the cerebral cortex may be defined by temporal or layer-specific expression of TFs (Arlotta *et al*., 2005; Molyneaux *et al*., 2007). While several studies have elaborated the importance of TFs in regulating cell-type specific dendrite development, the complex nature of the cellular and molecular mechanisms underlying this essential biological process are only beginning to be understood (Nanda et al., 2017; Puram and Bonni, 2013; Santiago and Bashaw, 2014; Yalgin *et al*., 2015; Sears and Broihier, 2016).

*Drosophila melanogaster* has proven to be an exceptionally powerful system for identifying and characterizing transcriptional targets and cellular pathways that operate as downstream effectors of TF-mediated dendrite morphogenesis. In particular, *Drosophila* multidendritic (md) sensory neurons have emerged as a powerful model for dissecting the molecular mechanisms underlying dendrite arbor specification and diversification, and studies using this model system have revealed numerous insights into the complex intrinsic and extrinsic regulatory mechanisms underlying cell-type specific dendrite development, including dendritic outgrowth, branching, maintenance, scaling, and tiling (Grueber, Jan and Jan, 2002; Jan and Jan, 2010; Tavosanis, 2021). These md neurons are categorized into four distinct morphological classes (Class I-IV; CI-CIV) based on their increasing orders of dendritic complexity (Grueber, Jan and Jan, 2002) thereby facilitating analyses of TF-mediated programs underlying dendritic morphological diversity.

Studies in *Drosophila* have begun to decode molecular mechanisms via which cell-type specific morphology TF networks converge on the cytoskeleton and other cellular pathways to drive dendrite arborization and spatial/functional compartmentalization (Jinushi-Nakao *et al*., 2007; Ye *et al*., 2007; Sulkowski *et al*., 2011; Ye *et al*., 2011; Iyer *et al*., 2012; Nagel *et al*., 2012; S.C. Iyer *et al*., 2013; Hattori *et al*., 2013; Ferreira *et al*., 2014; Yalgin *et al*., 2015; Sears and Broihier, 2016; Das *et al*., 2017; Clark *et al*., 2018; Yoong *et al*., 2020; Bhattacharjee *et al*., 2022), as well as initiator selector TF regulatory networks which regulate the expression of other TFs to govern a cascade of gene expression programs that drive neuronal differentiation (Ferreira *et al*., 2014; Corty, Tam and Grueber, 2016). Despite this significant progress, much remains to be discovered with respect to feed forward and reciprocal TF regulatory cascades, as well as the molecular mechanisms and cell biological processes by which TFs direct dendrite development through spatiotemporal modulation of cytoskeletal dynamics and other diverse signaling pathways. Moreover, the view that different TFs are dedicated to distinct phases of neuronal morphogenesis is likely an oversimplification, and recent studies have shown that TFs continue to play important regulatory roles in postmitotic neurons with respect to mediating distinct aspects of neuronal and dendritic development (Pai and Moore, 2021).

To identify and characterize morphology TFs that are required in specifying cell-type specific dendritic arborization, we previously reported on functional genomic analyses of two morphologically distinct md neuron subtypes, CI and CIV md neurons (E.P.R. Iyer *et al*., 2013). Cell-type specific CI vs. CIV transcriptomic profiling combined with a systematic loss-of-function genetic screen identified differentially enriched or depleted genes in CI vs. CIV md neurons, and in this study, we characterized a limited subset of 37 TFs that were differentially expressed (E.P.R. Iyer *et al*., 2013). In addition to differentially expressed TFs, we also conducted a separate genetic screen of TFs that were expressed in these neurons, albeit not differentially expressed, from which we identified a previously uncharacterized C2H2 zinc finger transcription factor encoded by the *Drosophila melanogaster CG3995* gene. Based on our pilot screen, *CG3995* was selected for further analyses as a result of the highly penetrant defects we observed on cell-type specific md neuron dendritic development. *CG3995* encodes an evolutionarily conserved zinc-finger BED-type (ZBED) protein containing a DNA binding domain found in chromatin boundary element binding proteins and transposases (Aravind, 2000). Members of the ZBED protein family in *Drosophila* include the proteins boundary element-associated factor of 32 kD (BEAF-32) and DNA replication-related element factor (Dref) which have been shown to antagonize each other in competing for binding to insulator or chromatin boundary elements (Aravind, 2000). Dysregulation of *CG3995* in md neurons leads to dendritic hypotrophy indicative of a functional role in directing proportional growth and branching. Due to the proportional reductions in both dendritic branching and overall dendrite growth, mutant neurons exhibit a “dwarfed” phenotype and thus we suggest this gene be named *bedwarfed* (*bdwf*) owing to the zinc-BED domain and the dendritic growth/branching defect leading to a stunted arborization pattern. Phenotypic analyses indicate Bdwf primarily regulates dendrite growth and branching via at least two distinct mechanisms in md neurons involving transcriptional and translational regulation. At the transcriptional level, we demonstrate that the Cut homeodomain transcription factor, which is known to exhibit cell-type specific expression and drive cell-type specific dendritic arborization in md neurons (Grueber *et al*., 2003), positively regulates Bdwf expression where this protein acts as a downstream effector of Cut-mediated dendrite morphogenesis, whereas Bdwf overexpression negatively regulates Cut expression revealing a reciprocal regulatory relationship between these TFs. At the translational level, proteomic analyses reveal that Bdwf primarily interacts with ribosomal proteins in CIV md neurons to regulate dendritic and cytoskeletal architecture. Further, our data indicate that Bdwf is required for proper trafficking of ribosomes along the dendritic arbor in both CI and CIV md neurons. Finally, we demonstrate that *bdwf* knockdown inhibits protein translation, suggesting that Bdwf regulates dendritogenesis by promoting global protein synthesis, perhaps through Bdwf interactions with ribosomal proteins. Collectively, these results provide new insights into complex combinatorial and multi-functional roles of TFs in determining cell-type specific dendrite development.

## Results

### *bdwf* dysregulation results in dendritic hypotrophy

The *bdwf* gene was identified in a RNAi-mediated phenotypic screen designed to identify predicted TF/DNA binding proteins that exhibit functional roles in regulating cell-type specific dendrite morphogenesis. The *bdwf* gene encodes a single mRNA isoform bearing two coding exons and predicted to produce a protein of 322 amino acids (37.5kDa) (Figure 1 A). Bdwf belongs to the C2H2 family of zinc-finger, BED-type (ZBED) transcription factors and bears N-terminal ZBED (amino acids 6-47) and Myb/SANT (amino acids 78-143) DNA binding/protein-protein interaction domains. To dissect the functional role(s) of *bdwf* in regulating cell-type specific dendritogenesis, we performed loss-of-function (LOF) and gain-of-function (GOF) analyses via gene-specific RNAi knockdown, mutant analysis or overexpression of full-length *bdwf* in multiple md neuron subclasses (CI; CIII; CIV). For mutation analysis, we a P{XP} transposon insertion in the *bdwf* 5’ region, *bdwf^d05488^* (Figure 1 A). Homozygous mutation for *bdwf^d05488^* displayed dendritic hypotrophy, including highly collapsed terminal branching and reduced field coverage in CIV md neurons (Figure S1 A, B, D, E). To verify the specificity of the *bdwf^d05488^* allele, we performed rescue experiments and found that the observed dendritic defects were fully rescued when a single copy of *UAS-bdwf-FLAG-HA* was expressed in CIV neurons in a homozygous *bdwf^d05488^* mutant background (Figure S1 A, C-E), thereby revealing a cell-autonomous requirement for *bdwf* in these neurons to promote dendritic arborization. To assess the putative role of *bdwf* in regulating cell-type specific dendritogenesis, we used the *GAL4-UAS* binary expression system for RNAi-mediated knockdown of *bdwf* in CI, CIII, or CIV md neurons. Two independent, gene-specific RNAi lines (*bdwf-IR*) with no predicted off-targets were tested and the transgenic line displaying stronger phenotypic penetrance was used for all subsequent RNAi analysis. CIV-specific *bdwf-IR* knockdown was highly penetrant and resulted in a significant dendritic hypotrophy manifesting as reductions in both dendritic branching and length in all three CIV md neuron subtypes (ddaC, v’ada, vdaB) (Figure 1B, C, K, L). Across the three CIV md neuron subtypes, *bdwf-IR* resulted in an average −26% reduction in total dendritic length (TDL), and an average - 27.7% reduction in total dendritic branches (TDBs) in CIV neurons relative to controls (Figure 1 B, C, Q). *bdwf* LOF phenotypes were most pronounced in CIII neurons. In contrast to the elaborate space-filling branching pattern of CIV neurons, CIII neurons are characterized by the presence of filopodial-like terminal branches (Figure 1B, E). When *bdwf-IR* was expressed under the control of a *CIII-GAL4* driver, we observed an average −48.5% reduction in TDL and −53% reduction in TDB (Figure 1 E, F, M, N, Q). *bdwf-IR* knockdown in CI neurons resulted in a similar degree of dendritic hypotrophy as observed in CIV neurons leading to an average reduction of −29.6% and −24.7% in TDB and TDL, respectively (Figure 1 H, I, O, P, Q).

**Figure 1:**
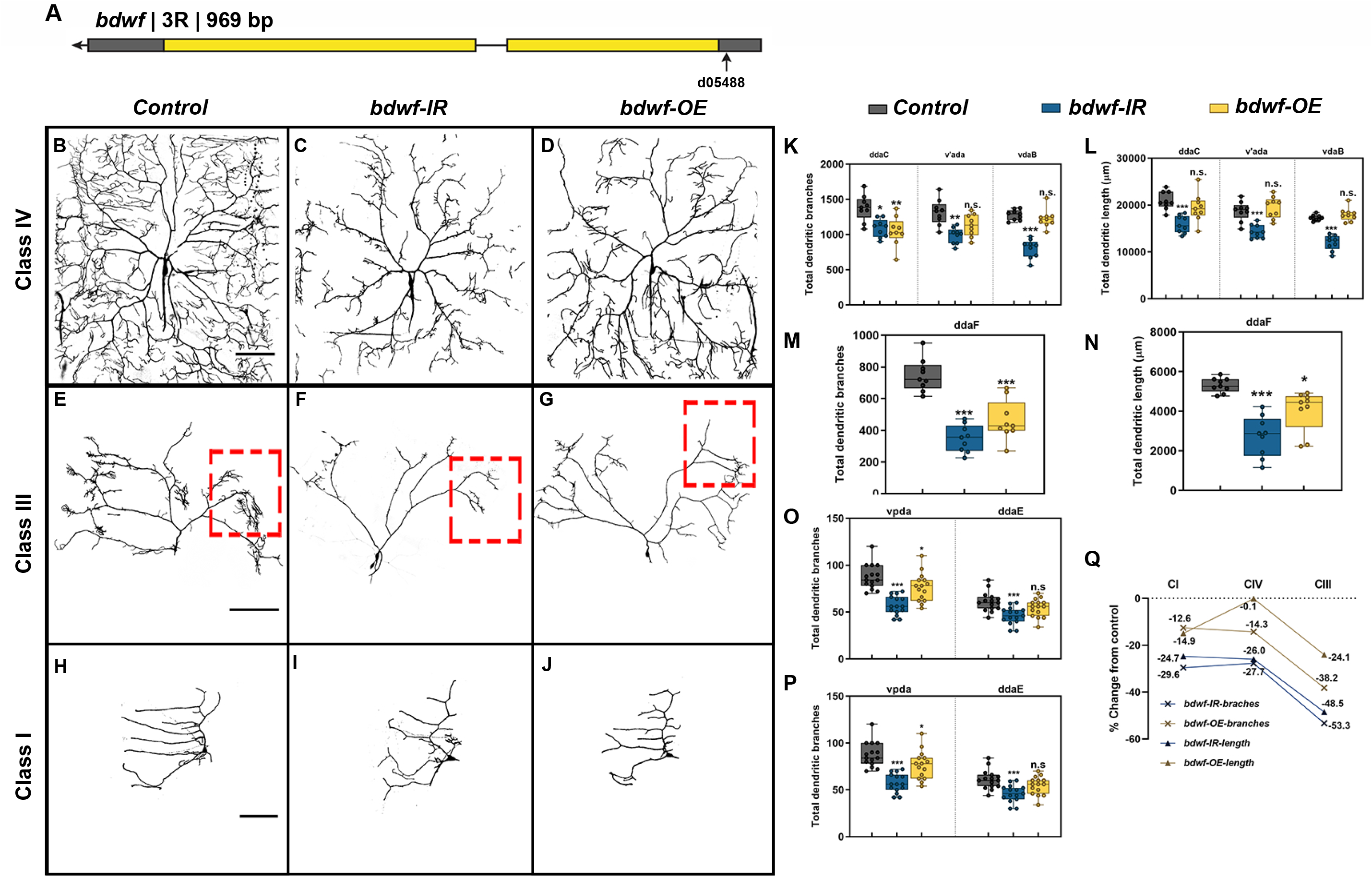
Optimal levels of Bdwf are essential for proper dendrite morphogenesis. (A) Schematic diagram showing the Bdwf gene and the location of the mutant allele used in the study. LOF and GOF results in strong and penetrant dendritic growth and branching defects in CIV (B-D), III (E-G) and I neurons (H-J). Scale bar, 50 μm. (K-P) Quantification of the total dendritic branches and total dendritic length are shown for CIV (K, L), CIII (M, N) and CI neurons (O, P). *p ≤ 0.05, **p ≤ 0.01, ***p ≤ 0.001 (One-way ANOVA with Dunnett’s multiple comparison test). Number of samples, n=14-15 for CI, n=9 for CIII and n=8-9 for CIV. (Q) Total dendritic length and total dendritic branches for each condition are represented as percentage change from control. Genotypes: (B-D, K, L, Q) *Gal4^477^, UAS-mCD8∷GFP/+; +/+* or *Gal4^477^, UAS-mCD8∷GFP/+; UAS-bdwf-IR/+* or *Gal4^477^, UAS-mCD8∷GFP/+; UAS-bdwf-FLAG-HA/+*. (E-G, M, N, Q) *GAL4^ppk^,GAL80^ppk^,UAS-mCD8∷GFP/+;+/+* or *GAL4^ppk^,GAL80^ppk^,UAS-mCD8∷GFP/+;UAS-bdwf-IR/+* or *GAL4^ppk^,GAL80^ppk^,UAS-mCD8∷GFP/+;UAS-bdwf-FLAG-HA/+*. (H-J, O, P, Q) *GAL4^221^,UAS-mCD8∷GFP/+ or GAL4^221^,UAS-mCD8∷GFP/UAS-bdwf-IR* or *GAL4^221^,UAS-mCD8∷GFP/UAS-bdwf-FLAG-HA*.

Since *bdwf* LOF resulted in dendritic hypotrophy, we next sought to investigate how *bdwf* overexpression (*bdwf-OE*) may impact dendritic morphogenesis using the same *UAS-bdwf-FLAG-HA* transgene used in rescue studies described above. *bdwf-*OE phenotypic analyses were conducted in CI, CIII, and CIV md neurons resulting in mild-to-severe dendritic hypotrophy as quantified by reduced TDL and/or TDB. CIII neurons displayed the strongest phenotypic defects with an average −38.2% reduction in TDB and −24.1% reduction in TDL (Figure 1 G, M, N, Q). On average, CI md neurons displayed a moderate reduction in TDL and TDB by −14.9% and −12.6%, respectively (Figure 1 H, O, P, Q). *bdwf* overexpression had the mildest effect on CIV neurons which displayed an average −14.3% reduction in TDB, albeit this defect was only observed in the dorsal most CIV md neuron subtype (ddaC), but showed no significant change (−0.1% reduction) in TDL relative to control (Figure 1 D, K, L, P). Collectively, LOF and GOF phenotypic analyses indicate that cell-type specific dendritic morphogenesis is sensitive to the Bdwf levels in CI, CIII, and CIV md neurons.

### *bdwf* regulates proportional dendritic growth and branching in morphologically divergent neurons

Qualitative analyses of *bdwf* LOF/GOF phenotypes revealed that *bdwf* LOF mutants appear to have “dwarfed” dendritic arbor morphology relative to control, manifesting as concomitant reductions in total dendritic growth and higher order branching across md neuron subclasses, whereas GOF analyses suggested that there were larger effects on higher order dendritic branching, most notably in the more morphologically complex CIII and CIV neurons (Figure 2). To quantify the subclass-specific effects of *bdwf* dysregulation on cell-type specific arborization patterns, we calculated dendritic branch density (DBD) by normalizing the TDB by TDL for each md neuron subtype. This metric is effectively the inverse of the average branch length, which is useful to distinguish between arbor size and complexity (Brown, Gillette, and Ascoli, 2008). In five of the six neuronal subtypes we analyzed, *bdwf-IR* knockdown displayed no statistically significant changes in branch density compared to controls (ddaC, v’ada, ddaF, ddaE, vpda). The only exception to this observation was the CIV vdaB ventral subtype, which showed a mild DBD reduction (−9.5%) (Figure 2 A, B, D-G). The observation that *bdwf* LOF mutants quantitatively resemble their WT control counterparts with respect to DBD indicates that the reduction in dendritic branching is highly proportional to the reductions in total dendritic length (Figure 2 G). These data suggest that Bdwf plays a role in regulating proper dendritic arborization by facilitating proportional dendritic growth and branching in morphologically divergent neurons. Conversely, quantitative analyses of *bdwf* GOF phenotypes revealed significant reductions in DBD for CIII and CIV neurons, whereas no significant change was observed in CI neurons (Figure 2 A, C-F). CIII neurons displayed the most reduction in branch density (−31.2%) followed by CIV neurons (−14.25%), whereas CI neurons displayed no significant change in DBD (+2.5%) relative to control (Figure 2G). This suggests that Bdwf overexpression exerts distinct effects on dendritic branching vs. growth in more morphologically complex CIII and CIV neurons as compared to simpler CI neurons, which may be due to cell-type specific differences in Bdwf interactors across md neuron subclasses.

**Figure 2:**
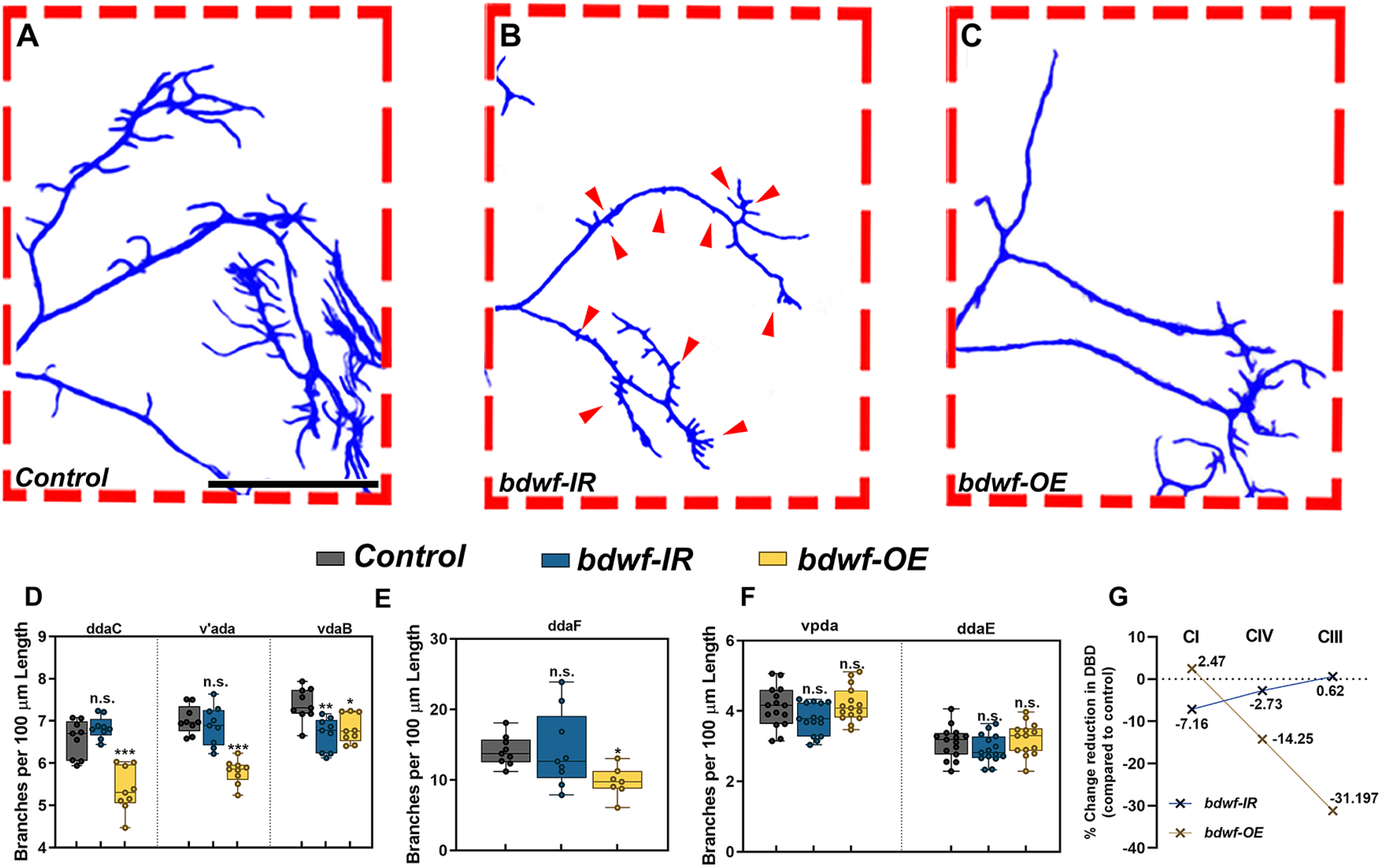
bdwf is required for proportional dendritic growth and branching. (A-C) Compared to wild-type (A), Bdwf LOF (B) results in a proportional reduction of TDB with respect to TDL, seen by the reduction in the size and numbers of CIII dendritic termini (red arrows). In contrast Bdwf GOF (C) results in suppression of dendritic termini. Scale bar, 100 μm. (D-F), Quantification of TDBD for CIV, III and I neurons (D, E and F) respectively. (G) Class-specific effects of Bdwf LOF and GOF on TDBD represented as percentage change from control. *p ≤ 0.05, **p ≤ 0.01, ***p ≤ 0.001 (One-way ANOVA with Dunnett’s multiple comparison test). n=14-15 for CI, n=7,9 for CIII and n=8,9 for CIV. Genotypes: *Gal4^ppk^, GAL80^ppk^, UAS-mCD8∷GFP/+; +/+* or *GAL4^ppk^,GAL80^ppk^,UAS-mCD8∷GFP/+;UAS-bdwf-IR/+* or *GAL4^ppk^,GAL80^ppk^,UAS-mCD8∷GFP/+;UAS-bdwf-FLAG-HA/+*(A-C, E, G). *GAL4^477^, UAS-mCD8∷GFP/+; +/+* or *Gal4^477^, UAS-mCD8∷GFP/+; UAS-bdwf-IR/+* or *Gal4^477^, UAS-mCD8∷GFP/+; UAS-bdwf-FLAG-HA/+* (D, G). *GAL4^221^,UAS-mCD8∷GFP/+ or GAL4^221^,UAS-mCD8∷GFP/UAS-bdwf-IR* or *GAL4^221^,UAS-mCD8∷GFP/UAS-bdwf-FLAG-HA* (F, G).

### Bdwf exhibits nucleocytoplasmic localization in md neurons

Given the LOF and GOF phenotypes observed with *bdwf* dysregulation, we sought to investigate the expression and subcellular localization of Bdwf protein in md neuron subclasses. Immunohistochemistry was performed on third instar larval filets revealing Bdwf protein expression in multiple md neuron subclasses. Bdwf exhibited punctate nucleocytoplasmic localization with qualitatively higher levels of cytoplasmic expression in control md neurons (Figure 3 A, B). In addition to Bdwf expression in md neuron subclasses, we also observed expression in adjacent epithelial and/or muscle cells between which the md neurons are sandwiched (Figure 3 A, B). To verify the specificity of the antibody and to document the efficacy of the *bdwf-IR* knockdown transgene, we analyzed Bdwf expression levels in control vs. *bdwf-IR* CIV md neurons, which revealed a significant reduction of Bdwf protein expression levels (Figure S1 F-H). In contrast to the more cytoplasmically distributed Bdwf puncta observed in control neurons, *bdwf* overexpression, using a pan-md *GAL4* driver, led to a dramatic shift to a predominantly nuclear subcellular localization for Bdwf expression (Figure 3 C, D). To dissect putative effects of Bdwf domains on subcellular localization, we generated structure-function mutant transgenes that express either the BED DNA binding domain alone (*UAS-bdwf-BED-HA*) or a version of *bdwf* lacking only the BED domain (*UAS-bdwf-ΔBED-myc*). Using a CIV-specific *GAL4* driver, we expressed these structure-function *bdwf* variants and immunostained for epitope tags associated with each transgene to visualize subcellular distribution. Expression studies revealed that the transgene lacking the Bdwf BED domain (*UAS-bdwf-ΔBED-myc*) was localized to the cytoplasm (Figure 3 E, F), whereas the transgene bearing only the Bdwf BED (*UAS-bdwf-BED-HA*) was predominantly localized to the nucleus (Figure 3 G, H). These data suggest that the Bdwf BED domain regulates nuclear localization.

**Figure 3:**
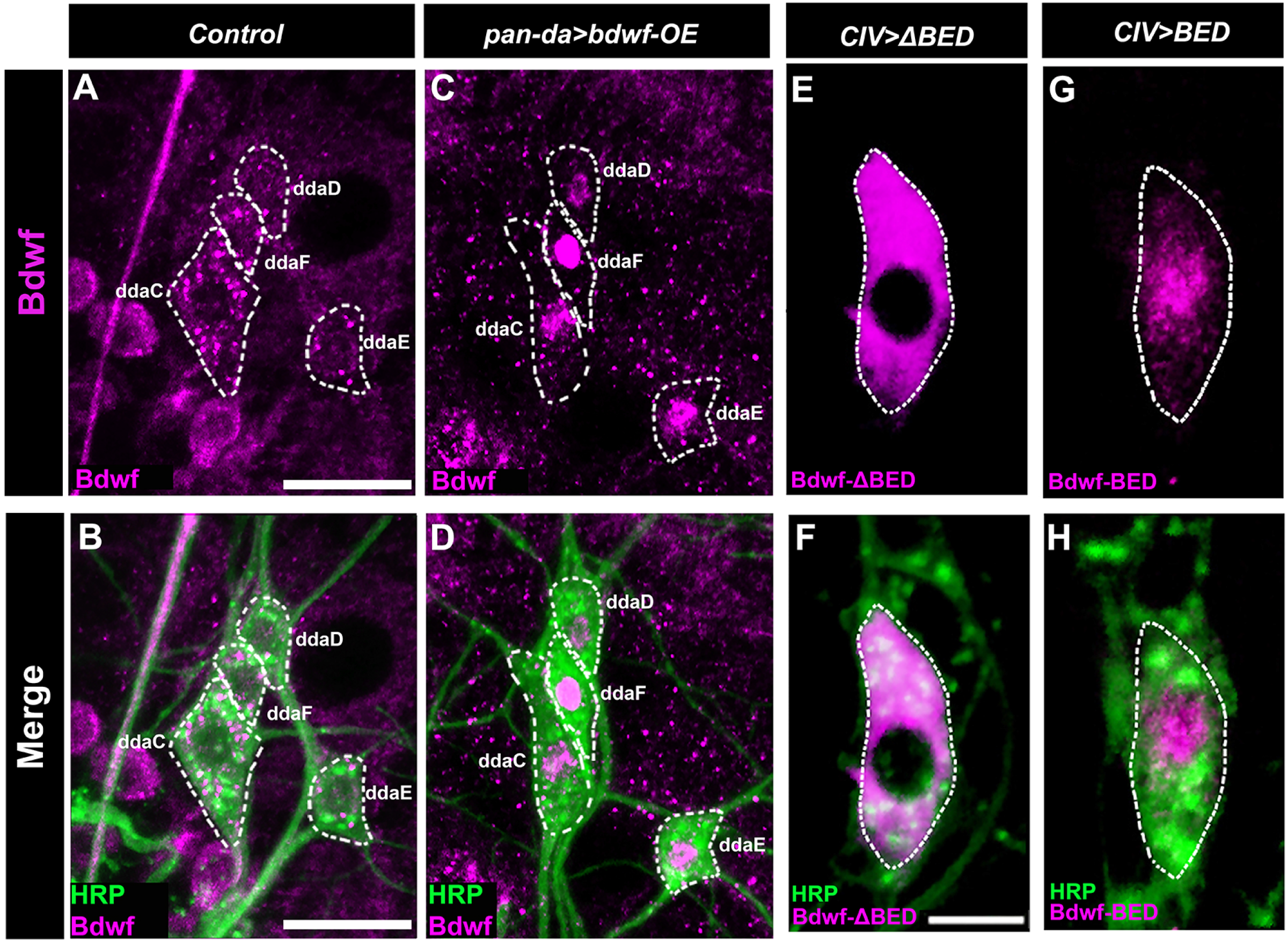
Bdwf exhibits nucleo-cytoplasmic localization. (A-D) Representative confocal image of WT and Bdwf overexpressing dorsal cluster da neurons labeled with HRP (green) and anti-Bdwf antibody (Magenta) to mark md neurons and Bdwf. md neuron cell bodies are highlighted by dotted white lines to represent Bdwf expression and localization in class I, III and IV neurons. In wild-type, Bdwf shows weak nuclear localization and strong punctate cytoplasmic expression (A, B). Overexpression of *bdwf* transgene with the pan-da *GAL4^217^* driver produces a dramatic increase in nuclear localization (C, D) as compared to wild-type, while still maintaining punctate expression in the cytoplasm. Note, that Bdwf is also expressed in puncta in surrounding epithelial and muscle tissue not labeled by HRP. Scale bar represents 20 μm. (E-H) BED domain is essential for Bdwf nuclearization. Representative confocal images of CIV md neurons expressing either *UAS-Bdwf-BED* or *UAS-Bdwf-ΔBED* using a class-IV specific driver. md neurons are immunolabeled with FITC-HRP (green) and anti-HA antibody (Magenta) to mark neurons and Bdwf mutant variants. Scale bar represents 5μm. Genotypes: *GAL4^217^/+;+/+* (A, B), *GAL4^217^/+; UAS-bdwf-FLAG-HA /+* (C, D), *GAL4^477^/UAS-ΔBED-myc;+/+* (E, F), *GAL4^477^/UAS-BED-HA;+/+* (G, H)

### Cut and Bdwf exhibit a reciprocal regulatory relationship with respect to expression

Previous studies have demonstrated that TF-mediated regulation of cell-type specific md neuron dendritic diversification is subject to TF regulatory networks whereby select TFs regulate the expression of other TFs to target downstream effectors that drive morphological diversity (Ferreira *et al*., 2014; Corty, Tam and Grueber, 2016; Pai and Moore, 2021). As LOF and GOF of analyses of *bdwf* revealed the strongest phenotypic defects in CIII md neurons, we hypothesized that Bdwf may operate in a TF regulatory network with the Cut (Ct) homeodomain transcription factor, which is expressed at highest levels in CIII neurons and is known to drive cell-type specific dendritic arborization in select md neuron subtypes (Classes II-IV) (Grueber *et al*., 2003). Moreover, there is a high degree of phenotypic similarity with respect to dendrite morphogenesis defects observed for *bdwf* and *ct* dysregulation. For example, previous studies have demonstrated that formation of the actin-rich terminal filopodial branches, characteristic of CIII neurons, is dependent upon Ct (Grueber *et al*., 2003), which are found to be significantly reduced in *bdwf* GOF mutants (Figure 2 E).

To explore a putative regulatory relationship between Ct and Bdwf, we first tested a hypothesis that Ct may positively regulate *bdwf* expression. To address this hypothesis, we isolated control CI md neurons and CI neurons in which we ectopically overexpressed Ct, and then performed qRT-PCR analyses to measure potential effects on *bdwf* mRNA expression levels. We observed that expression of a single copy of *ct* (*UAS-ct*) in CI neurons led to over a 4-fold increase in *bdwf* mRNA levels (Figure 4 A). In comparison, *ct* mRNA expression levels were found to be upregulated by ~14-fold with respect to control CI neurons (Figure 4 A). To assess how changes in mRNA levels may translate to protein levels, we ectopically overexpressed Ct in CI neurons and assayed for changes in native Bdwf protein levels via immunostaining. Ectopic Ct expression in CI neurons led to over 110% increase in Bdwf protein expression in these neurons relative to controls (Figure 4 B, D-K). In addition, we noted a mild, but significant, increase in nuclear Bdwf levels when Ct was ectopically expressed as quantified by nuclear to whole soma ratio of Bdwf expression (Figure 4 C). Expression analyses revealed Bdwf labeling in CI md neurons (Figure 3 A, Figure 4 D-G), suggesting that, at least in CI neurons, Ct is not required for Bdwf protein expression given that Ct is not normally expressed at detectable levels in these neurons (Grueber *et al*., 2003; Das *et al*., 2017). To examine whether this holds true for neurons that normally express Ct, we used mosaic analysis with a repressible cell marker (MARCM) to generate *ct^c145^* null mutant CIV neuron clones and stained for Bdwf. These analyses revealed that *ct* is not solely required for Bdwf protein expression in Ct-positive CIV neurons (Figure S2 A-C). Collectively, these analyses indicate that Cut exerts positive regulatory effects on *bdwf* mRNA and Bdwf protein expression in md neurons.

**Figure-4:**
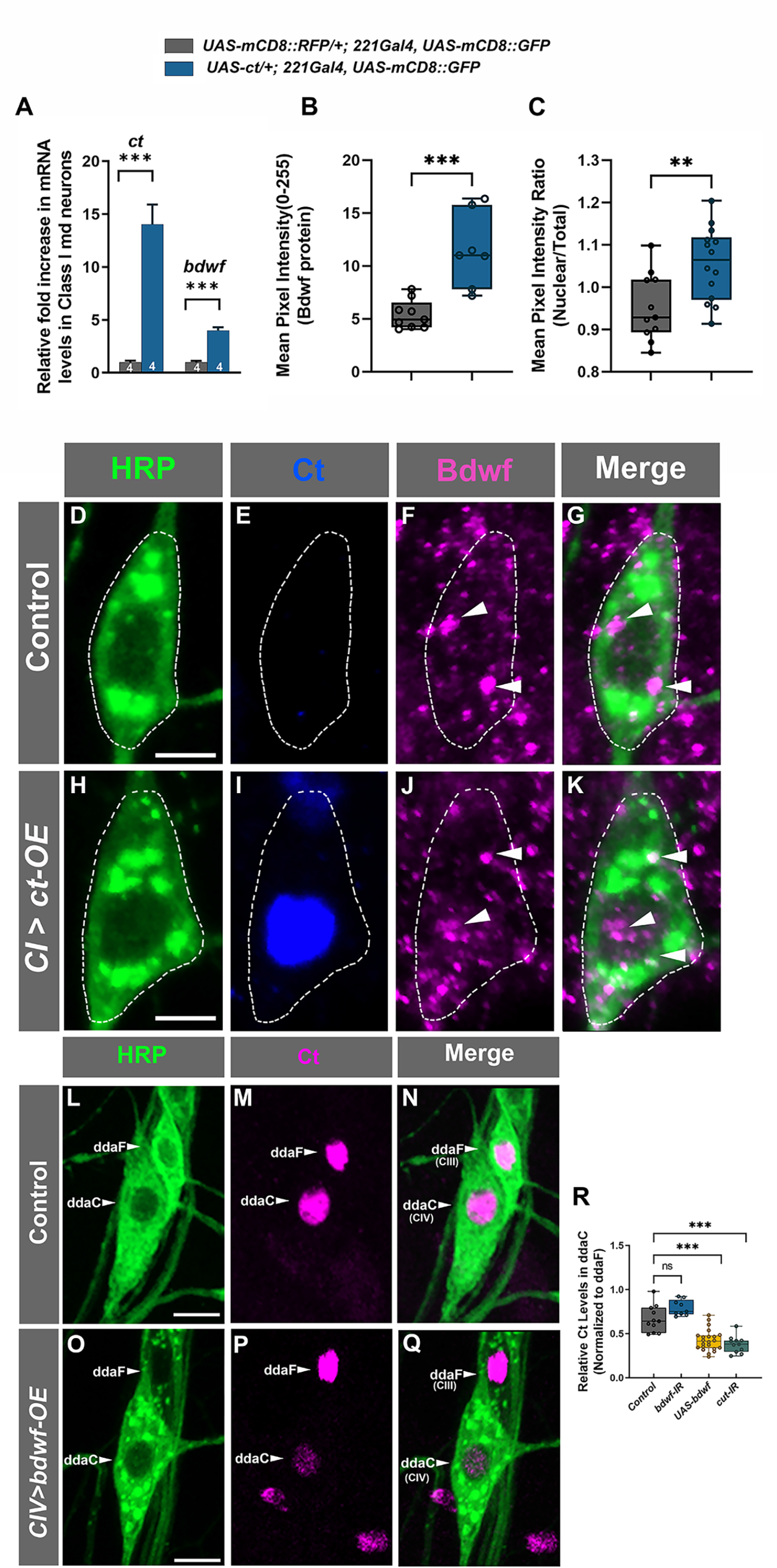
Cut and Bdwf reciprocally regulate one-another. (A-K) Cut positively regulates Bdwf expression. Immunohistochemical labeling of wild-type (D-G) and ectopically Cut expressing (H-K) ventral CI md neurons immunolabeled with HRP (Green), Cut (Blue) and Bdwf (Magenta) antibodies. (L-Q) Bdwf negatively regulates Cut expression. Compared to wild-type (L-N) Bdwf overexpression results in a strong and significant suppression of Cut immunostaining levels in CIV ddaC neurons (O-Q) immunolabeled with HRP (Green), Cut (Magenta). Scale bar, 5 μm. (A) RT-PCR quantification of relative mRNA levels of Cut and Bdwf in wild-type and Cut-overexpressing class-I neurons. (B) Mean pixel intensity quantification Bdwf immunostaining in wild-type and Cut expressing CI vpda neurons. (C) Nuclear-to-Cytoplasmic pixel intensity quantification of wild-type and Bdwf expressing CI vpda neurons showing increased nuclearization with Cut expression. (R) Mean pixel intensity of Cut in CIV ddaC neurons normalized to CIII ddaF signal. Error bars represent +/− SEM. **p ≤ 0.01, ***p ≤ 0.001(Unpaired Student’s T-test or One-way ANOVA with Dunnett’s multiple comparison test). The number of samples (n value) for (A) is represented is indicated by the number inside the bar, for (B), n=9 for WT and n=9 for *ct-OE*, for (C) n=11 for WT and 14 for *ct-OE*, for (R), n is between 9-22 per genotype. Genotypes: *UAS-cut; Gal4^221^, UAS-mCD8∷GFP/+* or *+/+; Gal4^221^, UAS-mCD8∷GFP/+* (A-K). *Gal4^477^, UAS-mCD8∷GFP/+* or *Gal4^477^/UAS-mCD8∷GFP /UAS- Bdwf -FLAG-HA* (L-Q), *Gal4^477^, UAS-mCD8∷GFP/+* or *Gal4^477^/UAS-mCD8∷GFP /UAS-Bdwf-FLAG-HA* or *Gal4^477^/UAS-mCD8∷GFP/+; UAS- Bdwf -IR/+* or *Gal4^477^/UAS-mCD8∷GFP /+*; *UAS-ct-IR/+* (R).

Given the positive feed forward regulatory relationship we observed between Ct and Bdwf, we next sought to investigate whether Bdwf may exert any reciprocal regulatory effects on Ct expression. Since Bdwf overexpression induced a reduction in dendritic branch density in Cut positive neurons (CIII, CIV neurons), we hypothesized that Bdwf and Cut may exhibit a reciprocal regulatory relationship. To test this, we overexpressed Bdwf in CIV neurons and quantified Cut immunostaining levels normalized to adjacent wild-type CIII neurons. Expression of a single copy of *ct-IR* in CIV neurons resulted in a strong suppression of Cut immunostaining levels by ~−40% (Figure 4 R). Bdwf overexpression in CIV neurons likewise resulted in strong suppression of Cut expression, to nearly the same levels as observed with *ct-IR* expression (−36.1%, Figure 4 L-R), indicating that Bdwf can negatively regulate Cut in a reciprocal feedback loop when overexpressed in these neurons. To determine whether Bdwf may be required for Ct expression, we specifically knocked down *bdwf* in CIV neurons, which resulted in a 19% increase in Ct immunostaining when compared to controls; however, this change, while trending (p=0.057), did not rise to the level of statistical significance (Figure 4 R). Taken together, these data suggest that the Bdwf overexpression-induced dendritic hypotrophy phenotype is likely due, at least in part, to the negative feedback effects on Ct expression.

### Bdwf functions as a downstream effector of Cut-mediated dendritic arborization

Cut has previously been shown to induce dramatic dendritic hypertrophy in a dose-dependent manner in CI neurons (Grueber *et al*., 2003). To explore the functional consequence of Cut-Bdwf regulatory relationship, we investigated if Cut-induced dendritic complexity occurs via a Bdwf-dependent pathway. We hypothesized that if Bdwf acts as a downstream effector of Cut-mediated dendritic development, then disrupting *bdwf* in a genetic background sensitized by ectopic Ct overexpression should lead to a suppression of Cut-mediated dendritic hypertrophy. To test this hypothesis, we generated a line that stably expressed *UAS-cut* under the control of the CI md neuron driver *GAL4^221^* which was outcrossed to either *UAS-bdwf-IR*, *UAS-bdwf*, *UAS-cut* or *cut-IR* to assay the respective phenotypic effects on CI dendrite morphogenesis. To control for *GAL4-UAS* titration, we expressed *UAS-mCD8∷RFP* in the Cut overexpression control background. As expected, ectopic expression of Cut in CI neurons results in dramatic dendritic hypertrophy, which manifests as a sharp increase in lower order dendritic branch extension as well as the *de novo* formation of numerous terminal dendritic filopodia-like protrusions, relative to wild-type controls (Figure 5 A, B, E, F) (Grueber *et al*., 2003). Expression of *UAS-cut* (+ *UAS-mCD8∷RFP*) in CI vpda neurons resulted in over 7.5 fold increase in TDB and over 3 fold increase TDL with respect to wild-type controls (Figure 5 D, E, F). We first validated the effectiveness of this system by expressing *cut-IR*, which resulted in strong suppression of dendritic hypertrophy (Figure 5 C, E, F). Co-expression of *UAS-cut* and *UAS-bdwf-IR* likewise resulted in suppression of Cut-mediated dendritic hypertrophy with respect to dendritic growth and branching (Figure 5 D, E, F). These results indicate that Bdwf is required for Cut-induced dendritic hypertrophy in CI neurons.

**Figure 5:**
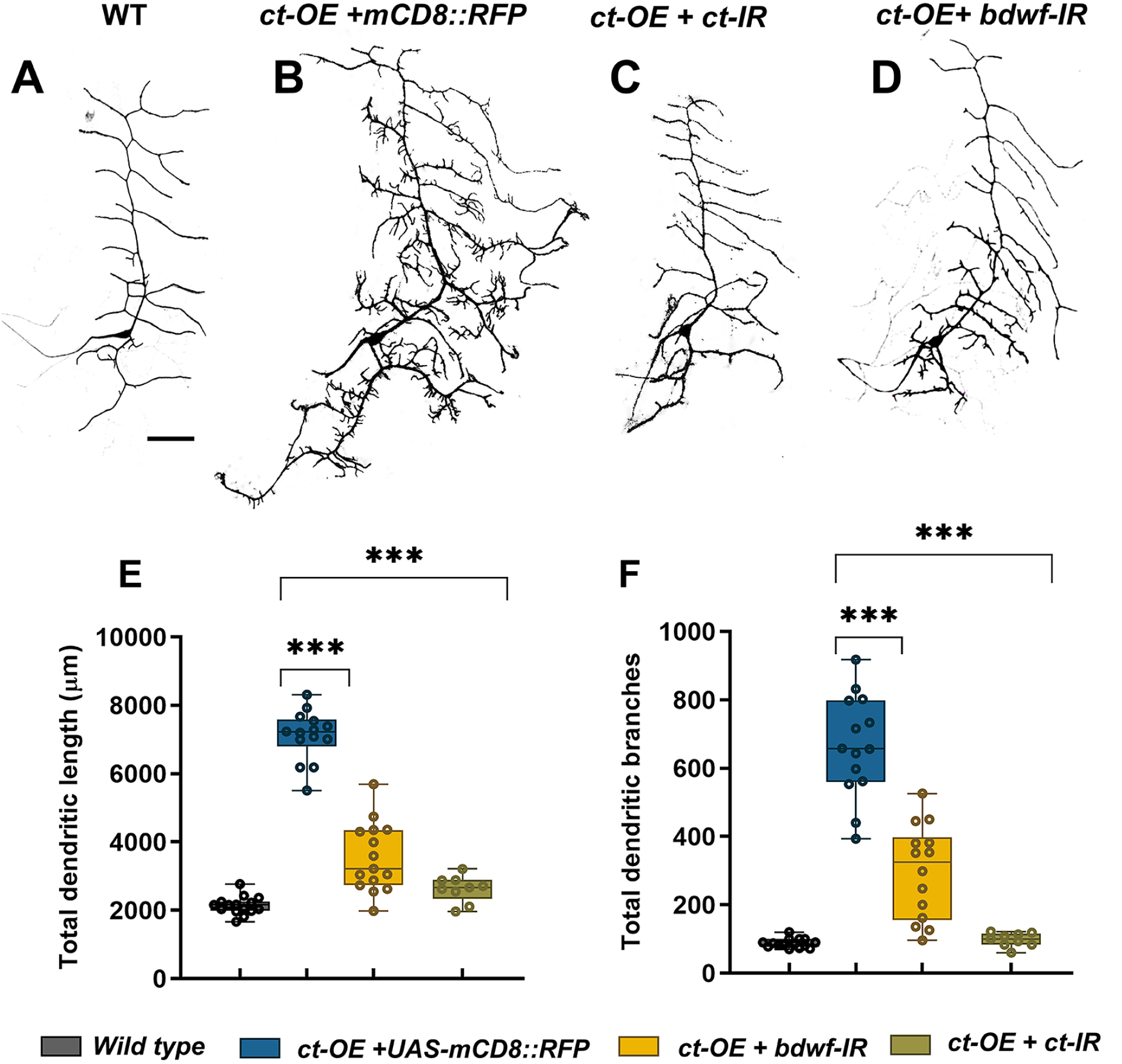
Bdwf is required for Cut-induced dendritic development. (A-D) Suppression of Cut GOF phenotype in CI neuron by Bdwf LOF. Representative images of wild-type (A), Cut-overexpressing (B), Cut-overexpressing with Cut-LOF (C), and Cut overexpressing with Bdwf–LOF (D) CI vpda neurons. Scale bar, 50 μm. Quantification of TDL (E), TDB (F), are shown. ***p ≤0.001(One-way ANOVA with Dunnett’s test for multiple comparison or Kruskal-Wallis with Dunn’s multiple correction test). n=15 for WT, n= 14 for *ct-OE+UAS-mCD8∷RFP*, n=14 for *ct-OE+bdwf-IR*, n=9 for *ct-OE+ct-IR*. Genotypes: *Gal4^221^, UAS-mCD8∷GFP/+* (A,E, F), *UAS-ct/+; Gal4^221^, UAS-mCD8∷GFP/+* (B,E, F), *UAS-Cut/+; Gal4221, UAS-mCD8∷GFP/UAS-ct-IR* (C,E, F), *UAS-ct/+; Gal4^221^, UAS-mCD8∷GFP/UAS- bdwf -IR* (D,E, F).

Given that Cut is not normally expressed in CI md neurons, we sought to test whether Bdwf overexpression could rescue dendritic hypotrophy defects observed when *ct* is knocked down in CIV md neurons, which normally express Ct protein. Relative to controls, CIV-targeted expression of *ct* RNAi (*ct-IR*) leads to significant reductions in both total dendritic length and number of dendritic branches (Figure S2 D, E, G, H). When we simultaneously knocked down *ct* (*ct-IR*) and overexpressed *bdwf* (*UAS-bdwf*) in CIV neurons, we observed a partial rescue resulting in a significant recovery of the reductions we observed for total dendritic length and number of dendrite branches (Figure S2 F, G, H). While the rescue did not achieve a full recovery of *ct-*mediated dendritic hypotrophy back to control morphology, this was predicted as numerous genes have been previously identified as downstream effectors of *ct-*mediated dendritic development (Nanda *et al*., 2017; Jinushi-Nakao *et al*., 2007; Sulkowski *et al*., 2011; Iyer *et al*., 2012; Nagel *et al*., 2012; S.C. Iyer *et al*., 2013; Ferreira *et al*., 2014; Das *et al*., 2017; Clark *et al*., 2018; Bhattacharjee *et al*., 2022). Collectively, these analyses reveal that Bdwf functions as a downstream effector Cut-mediated dendritic arborization.

### Bdwf interacts and colocalizes with ribosomal proteins

Endogenous Bdwf protein expression revealed strong cytoplasmic labeling and thus we sought to investigate potential mechanisms by which Bdwf may operate in the cytoplasm to regulate dendrite morphogenesis. To this end, we conducted affinity purification of Bdwf interacting proteins in combination with mass spectroscopy (MS) analyses on age-matched larval tissue. In addition to purifying Bdwf itself, the MS analyses identified over 70 proteins interactors of Bdwf, almost half of which were found to be ribosomal proteins (~44%) (Figure 6 A; Table S1). The gene ontology (GO) term “cytosolic ribosome” was found to be the most statistically enriched category in this dataset (~30-fold enrichment, p = 2.08E-42) (Table S1). Furthermore, co-immunostaining of Bdwf and one of the protein interactors, ribosomal protein S6 (RpS6), revealed colocalization of these proteins in the cytosol of CIV md neurons (Figure 6 B). These data reveal Bdwf physically interacts and colocalizes with ribosomal proteins.

**Figure 6:**
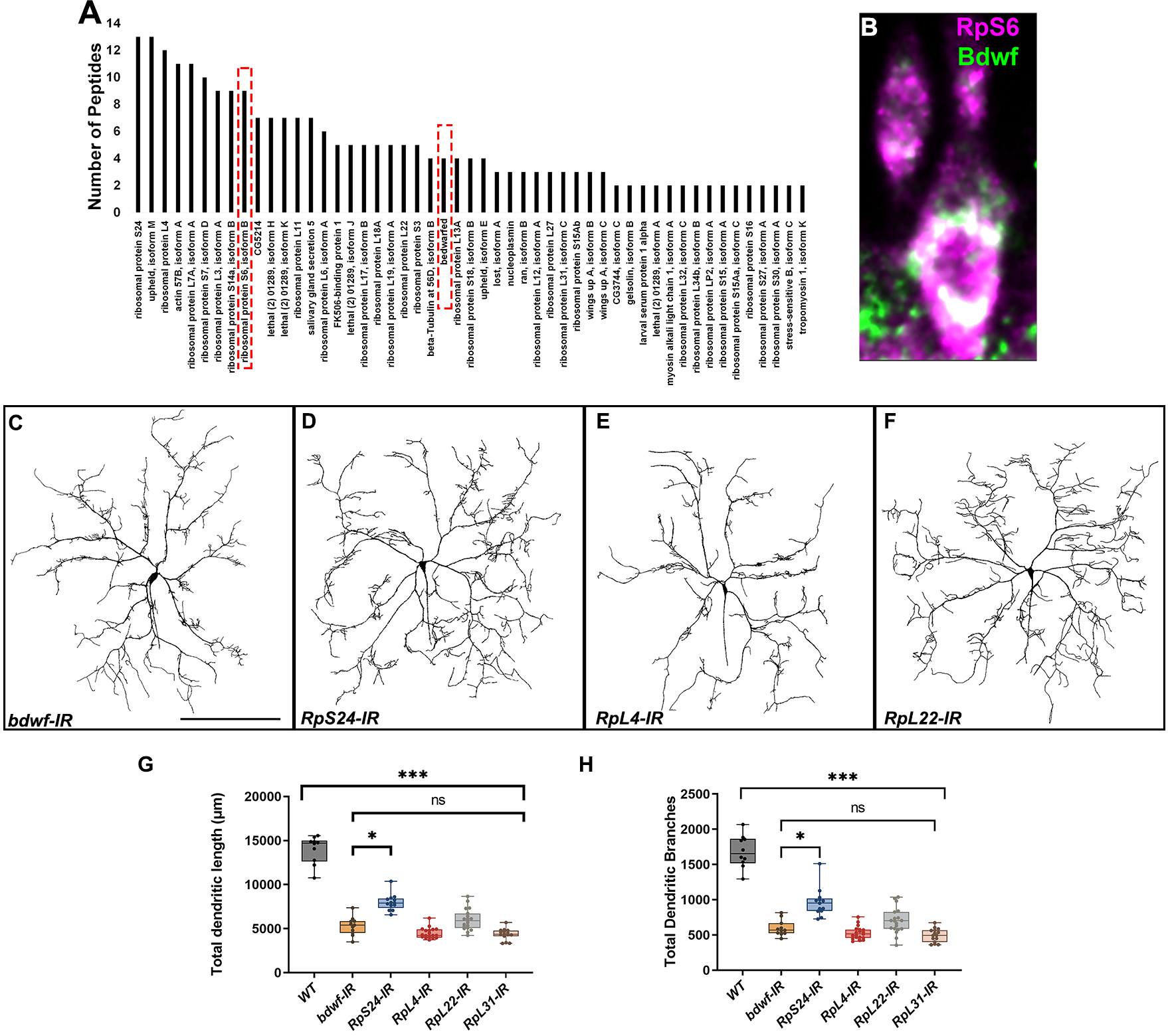
Bdwf interacts and colocalizes with ribosomal proteins. (A) Bdwf-interacting proteins identified from MS analysis of affinity purified Bdwf protein using our rabbit anti-Bdwf polyclonal antibody from homogenized third-instar larvae expressing *UAS-bdwf-FLAG-HA* protein driven by *GAL4^Act5C^* driver. Bdwf and ribosomal protein S6 are marked in dotted red box. Bdwf co-localizes with the 40s ribosomal protein S6 to a high degree in md neurons (B). Knockdown of ribosomal proteins lead to a significant reduction in the TDL (G) and TDB (H) compared to control. With the exception of *RpS24-IR*, knockdown of all the ribosomal proteins phenocopied bdwf knockdown with a high degree of penetrance. Representative images of *bdwf-IR*, *RpS24-IR*, *RpL4-IR*, and *RpL22-IR* (C-F). Scale bar, 200 μm Quantification of TDL and TDB in CIV ddaC neurons (G, H). * p≤0.05, *** p≤0.01. n=8 for WT, n=11 for *bdwf-IR*, n=13 for *RpS24-IR*, n=19 for *RpL4-IR*, n=19 for *RpL22-IR*, n=15 for *RpL31-IR*. Genotypes: *GAL4^477^,UAS-mCD8∷GDP/+;GAL4^ppk1.9^,UAS-mCD8∷GFP/+*, or *GAL4^477^,UAS-mCD8∷GDP/+;GAL4^ppk1.9^,UAS-mCD8∷GFP/UAS-bdwf-IR*, or *GAL4^477^,UAS-mCD8∷GDP/UAS-RpS24-IR;GAL4^ppk1.9^,UAS-mCD8∷GFP/+*, or *GAL4^477^,UAS-mCD8∷GDP/UAS-RpL4-IR;GAL4^ppk1.9^,UAS-mCD8∷GFP/+*, *GAL4^477^,UAS-mCD8∷GDP/+;GAL4^ppk1.9^,UAS-mCD8∷GFP/UAS-RpL22-IR*, or *GAL4*^477^,*UAS-mCD8∷GDP/UAS-RpL31-IR;GAL4^ppk1.9^,UAS-mCD8∷GFP/+*

We reasoned that if Bdwf interactors were functionally important for dendritogenesis, disruption of members of this protein complex should at least partially phenocopy *bdwf* LOF disruptions. To functionally validate the role of Bdwf interactors in dendritogenesis, we selected genes encoding both large and small subunit ribosomal proteins that were enriched in our MS dataset and performed gene-specific RNAi analyses in CIV neurons. CIV-targeted RNAi against *RpS24, RpL4, RpL22* and *RpL31* resulted in dendritic hypotrophy consistent with the defects observed with *bdwf* RNAi knockdown, including significant reductions in TDL and TDB relative to wild-type controls (Figure 6 C-F). In addition, with the exception of *RpS24-IR*, the reduction in the TDL and TDB for the ribosomal protein knockdowns were not significant when compared to *bdwf* RNAi (Figure 6 C-F, G, H). These results suggest that Bdwf might likely be functioning in conjunction with multi-protein ribosomal complexes to promote dendritic arborization.

Cell-type specific dendritic architecture is mediated by the organization and dynamics of cytoskeletal fibers (Das *et al*., 2017; Das *et al*., 2021; Kilo *et al*., 2021; Bhattacharjee *et al*., 2022). In the case of *Drosophila* md neurons, previous studies have demonstrated that the local distribution and organization of microtubule (MT) and F-actin fibers is sufficient in constraining arbor morphology (Nanda *et al*., 2020). Given that LOF for *bdwf* and ribosomal protein encoding genes produced similar defects in dendritic morphogenesis, we next examined how disruption of these genes may impact the underlying dendritic cytoskeleton. We investigated MT and F-actin cytoskeletal fibers in CIV neurons using multi-fluorescent reporter lines that label F-actin (*UAS-GMA∷GFP*) or MTs (*UAS-mCherry∷Jupiter*) (Das *et al*., 2017). The F-actin and MT intensities were then quantified by multichannel digital reconstructions as previously described (Nanda *et al*., 2021). CIV knockdown of *bdwf* or genes encoding ribosomal proteins severely diminished MT signal. In the case of *bdwf-IR*, a percentage change from control analysis revealed that, at 20 μm from the soma, there was ~50% reduction in MT signal which kept decreasing on dendrites distal to the soma with a nearly complete loss in MT signal at ~400 μm from the soma (Figure 7 A’, A’’, C’, C’’, H). With the exception of *RpS24*, knockdown of *RpL4, RpL22*, and *RpL31* ribosomal proteins had a similar effect on MT with nearly 70% reduction in MT signal at 20 μm from the soma which progressively decreased distal to the soma (Figure 7 E’, E’’, F’, F’’, G’, G’’, H). *RpS24* knockdown had a less severe effect on MT signal, with only ~40% reduction in MT signal at 20 μm from the soma. At the farthest distance from the cell body, 540 μm, the MT signal was reduced by ~77% (Figure 7 D’, D’’, H). In contrast to MTs, *bdwf-IR* led to a redistribution of the F-actin signal. Percent change from control analysis showed that at ~60-160 μm from the soma there was over 20% increase in F-actin signal, however beyond this distance from soma the F-actin signal continued to decrease on arbor most distal from the soma (Figure 7 A, C, I). Similar to their effect on MTs, knockdown of *RpS24, RpL4, RpL22*, and *RpL31* ribosomal proteins led to a decrease in the F-actin signal as a function of distance from the soma (Figure 7 D, E, F, G, I). Previous studies from our lab have shown that Ct can regulate effector molecules that target the cytoskeleton which in turn impacts dendritic morphogenesis (Das *et al*., 2017; Bhattacharjee *et al*., 2022). Moreover, we have also shown that Ct regulates the expression of ribosomal proteins to modulate dendritic complexity (Das *et al*., 2017). Since our study establishes that Cut and Bdwf have a reciprocal relationship, we examined defects associated with cytoskeletal architecture upon *ct* knockdown in CIV neurons, which revealed severely reduced F-actin labeling throughout the dendritic arbor and progressively diminishing MT signal as a function of distance from soma (Figure 7 B, B’, B’’, H, I). Taken together, our results indicate that *ct*, *bdwf* and genes encoding ribosomal proteins are required to support dendritic cytoskeletal organization and stability.

**Figure 7:**
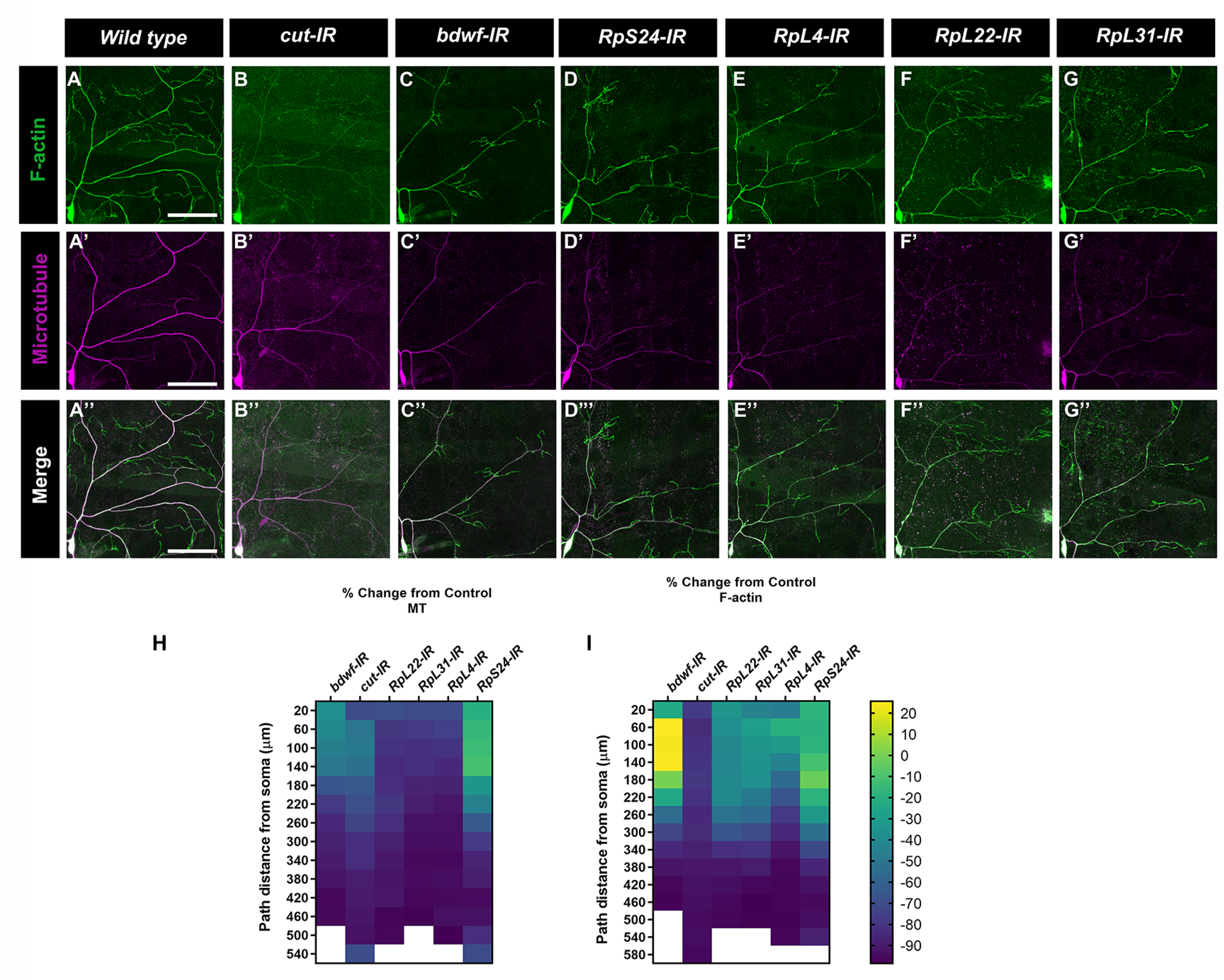
Bdwf and Cut interact with ribosomal proteins to modulate the cytoskeletal architecture. (A-G”) Representative images of CIV ddaC neurons labelled with CIV specific *UAS-GMA* (F-actin) and *UAS-mCherry∷Jupiter* (MT). (A-A”) Control and (B-G”) gene specific RNAi knockdown. (H, I). Heatmap showing the percent change from control of MT signal (H) and F-actin signal (I) as a function of the path distance from soma in knockdown conditions. n=8 for *bdwf-IR*, n=10 for *ct-IR*, n=10 for *RpL22-IR*, n=9 for *RpL31-IR*, n=10 for *RpL4-IR*, n=14 for *RpS24-IR*. Scale bar, 50 μm. Genotypes: *UAS-GMA;GAL4^477^,UAS-mCherry∷JUPITER/+;+/+* (A-A”), or *UAS-GMA;GAL4^477^,UAS-mCherry∷JUPITER/+; UAS-bdwf-IR/+* (C-C”, H, I), or *UAS-GMA;GAL4^477^,UAS-mCherry∷JUPITER/+;UAS-ct-IR/+*(B-B”, H, I), or *UAS-GMA;GAL4^477^,UAS-mCherry∷JUPITER/UAS-RpS24-IR;+/+* (D-D”, H, I), or *UAS-GMA;GAL4^477^,UAS-mCherry∷JUPITER/UAS-RpL4-IR;+/+* (E-E”, H, I), or *UAS-GMA;GAL4^477^,UAS-mCherry∷JUPITER/+;UAS-RpL22-IR/+* (F-F”, H, I), or *UAS-GMA;GAL4^477^,UAS-mCherry∷JUPITER/UAS-RpL31-IR;+/+* (G-G”, H, I).

### Bdwf is required for ribosomal trafficking and protein translation along the dendritic arbor

A previous study in *C. elegans* has shown that disruption of MTs altered ribosome localization in axons (Noma *et al*., 2017). Our cytoskeletal analyses showed that knockdown of *bdwf* severely disrupted MT stability along the dendritic arbor. We therefore sought to determine the effect of *bdwf-IR* on ribosome trafficking along the dendritic arbor. We expressed GFP tagged RpL10Ab (Thomas *et al*., 2012) under the control of either CI or CIV *GAL4* drivers. This GFP-tagged RpL10Ab is reported to be incorporated in both polysomes and monosomes (Thomas *et al*., 2012) and we used it as a proxy to investigate ribosome localization along the dendrites. In both CI and CIV control neurons, ribosome signal is found distributed along the arbor and at dendritic branch points (Figure 8 A, A’, E, E’). *bdwf* knockdown severely impaired ribosome trafficking along the arbor (Figure 8 B, B’, F, F’). Quantitative analyses revealed that in both CI and CIV neurons there was a significant reduction in ribosome density along the dendritic arbor (Figure 8 C, G) as well as in the number of branch points with ribosomes (Figure 8 D, H) in *bdwf-IR* compared to control. This suggests that Bdwf is required for ribosomal trafficking on dendritic arbors in these neurons.

**Figure 8:**
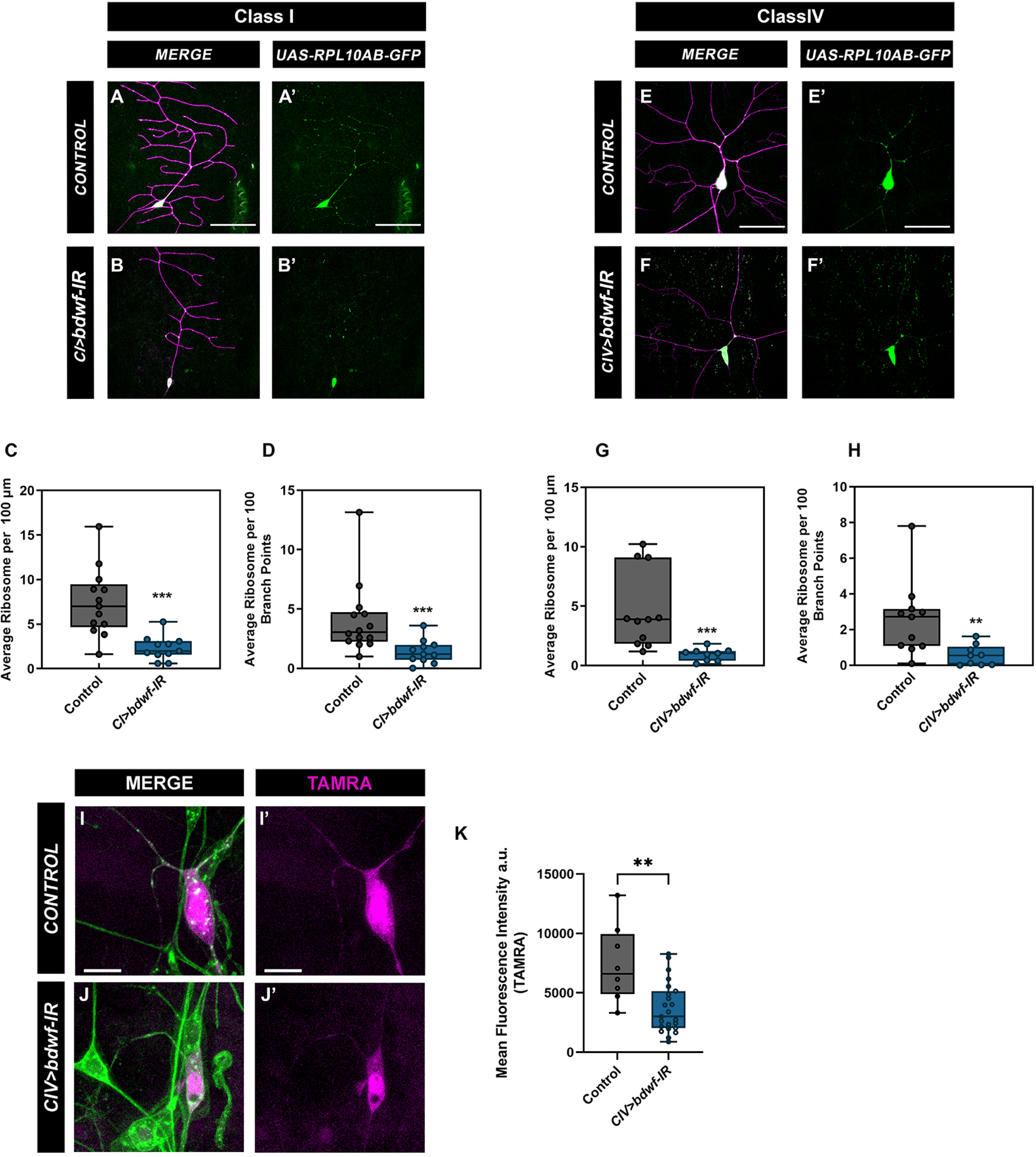
Bdwf is required for ribosome trafficking along the dendrite and protein translation. Knockdown of *bdwf* shows that Bdwf is required for ribosome trafficking along the dendritic arbor in both CI and CIV md neurons (A-H). (A, A’,B, B’, E, E’, G, and G’) Representative confocal image of WT and *bdwf-IR* CI and CIV md neurons respectively showing ribosome signal along the dendritic arbor. (C, D, G, H) Quantification of ribosome signal along the dendritic arbor normalized to length (C, G), or ribosome signal at branch points (D, H). Scale bar, 20 μm. Loss of Bdwf leads to disruption in protein translation (I-J). (I,I’J,J’) Representative images of control and *bdwf-IR* expressing CIV neurons labelled with TAMRA (magenta) to show translated protein and dendrites labelled with HRP (green). Scale bar represents 10 μm. **p < 0.01, ***p < 0.001(Student’s T-test or Mann-Whitney test), n= 14 for CI control (C, D), n=11 for CI *bdwf-IR* (C, D), n=11 for CIV control (G, H), n=9 for CIV *bdwf-IR* (G, H), n=8 for CIV control and n=23 for CIV *bdwf-IR* (K). Genotypes: *UAS-RPL10Ab∷GFP/+;GAL4^221^,UAS-mCD8∷RFP/+* (A, A’), *UAS-RPL10Ab∷GFP/+;GAL4^221^,UAS-mCD8∷RFP/UAS-bdwf-IR* (B, B”), *UAS-RPL10Ab∷GFP/+;GAL4^ppk^,UAS-mCD8∷RFP/+* (E, E’), *UAS-RPL10Ab∷GFP/+;GAL4^ppk^,UAS-mCD8∷RFP/UAS-bdwf-IR* (F, F’), *UAS-dMetRS^L624G^-3xmyc/+;GAL4^ppk^/+* (I, I’), *UAS-dMetRS^L624G^-3xmyc/+;GAL4^ppk^/UAS-bdwf-IR* (J, J’).

Our data demonstrate that Bdwf co-localizes with ribosomal proteins and is also required for ribosome localization along the dendritic arbor. We next wanted to determine the effect of *bdwf* knockdown on global protein translation. We expressed the mutant methionyl tRNA-synthetase (*UAS-dMetRS^L624G^-3xmyc*) under the control of a CIV *GAL4* in control and *bdwf-IR* conditions. MetRS^L274G^ causes methionyl-tRNA to be charged with azidonorleucine (ANL), allowing its incorporation into proteins which can then be used to visualize translated proteins through biorthogonal click chemistry labeling (Erdmann *et al*., 2015). In control neurons, *de novo* proteins tagged with the red-fluorescent dye tetramethyrhodamine (TAMRA) can be seen both in the cell body and along the dendrites (Figure 8 I, I’). In *bdwf-IR* animals, the TAMRA signal was restricted to the cell body with no discernable signal in the dendrites (Figure 8 J, J’). Quantitative analysis showed a significant reduction in the levels of translated protein in *bdwf-IR* compared to controls (Figure 8 K). Thus, collectively, our data reveals that Bdwf is required for ribosome trafficking along the dendritic arbor and for protein translation in md neurons.

## Discussion

Here we report on the characterization of a novel zinc-BED domain containing transcription factor, Bdwf, and its roles in regulating cell-type specific dendritogenesis in the *Drosophila* sensory neurons. In all three neuronal subtypes analyzed (CI, CIII, CIV md neurons), loss of *bdwf* led to dendritic hypotrophy and analyses indicate that Bdwf is required for proportional dendritic growth and branching. We present evidence to show that Bdwf exhibits nucleocytoplasmic localization and that Bdwf acts via at least two independent pathways to regulate dendritogenesis. In the complex CIII and CIV neurons, Bdwf and Cut exhibit a reciprocal regulatory relationship which promotes dendritic arborization in CIV neurons. Both Bdwf and Cut are required for MT and F-actin cytoskeletal organization and/or stability, and subsequent analyses reveal Bdwf associates with ribosomal proteins and is required for proper localization of the ribosome complex along the dendritic arbor and in turn protein synthesis. Position-Specific Iterated BLAST (PSI-BLAST) analysis of Bdwf protein sequence revealed that Bdwf shares sequence similarities with MSANTD4 and Zbed4 in mouse and ZBED4 in humans. While MSANTD4 has been linked to Huntington’s disease, and recent studies have linked ZBED4 to schizophrenia and Phelan-McDermid syndrome, little is known about the underlying mechanism of these proteins (Nyegaard *et al*., 2010; Mitz *et al*., 2018; Schenkel *et al*., 2021; Seefelder and Kochanek, 2021). Our study may provide new and valuable entry-points for further research on the role of these proteins in the nervous system function and dysfunction.

Our results indicate that the BED domain aids in Bdwf nuclearization, as its deletion leads to a predominantly cytoplasmic signal. One possibility is that Bdwf may self-associate to form a multimeric complex which might aid in its nuclearization. Studies of ZBED4/KIAA0636 revealed self-association via a hATC domain which was essential for its nuclear accumulation and DNA binding (Yamashita *et al*., 2007); however, sequence analyses of Bdwf did not identify a hATC domain, leaving open the possibility that, if Bdwf does rely on self-association for nuclear localization, this must occur via a dissimilar, and as yet unknown, mechanism. Beyond sensory neuron expression, which is the focus of this study, we also found Bdwf to be expressed in a diverse set of tissues including central nervous system neurons, muscle, epithelia, and ovaries (data not shown). Similar to our observation, the mouse homolog of Bdwf, ZBED4 has been reported to be expressed at different levels in many mouse tissues including brain, heart, ovary, and retina, and has both a nuclear and cytoplasmic localization in retina cones in both mouse and human (Farber *et al*., 2010; Saghizadeh *et al*., 2011).

For neurons to effectively receive and propagate signals, it is essential that they cover their receptive fields and form synaptic connections. As an organism grows, the organs and tissues also increase in size. Tissues keep up with this increasing size by two mechanisms, either by increasing the number of cells while keeping the size of the individual cell constant, or as in the case of certain neurons, such as the Purkinje cells, the axons and dendrites grow relative to the increase in body size (Wittenberg and Wang, 2007). With respect to dendritic arbor scaling, studies have demonstrated that animals grown under mild starvation conditions exhibit a decrease in arbor size that was proportional to the decrease in body size resulting in a miniaturized dendritic arbor (Shimono *et al*., 2014). In the case of miniaturized dendritic arbors, there are decreases in arbor size and total dendritic length, but often unaltered terminal branching, resulting in a net increase in dendritic branch density. In contrast, neurons with aberrant Insulin/IGF signaling or TORC1 signaling result in hypotrophic, dwarfed or simplified dendritic arbors that are distinguished from miniaturized arbors observed under starvation conditions in that there are proportional decreases in arbor size, terminal branching, and total length leading to dendritic branch densities that are not significantly different from control neurons (Shimono *et al*., 2014). Studies have shown that both cell autonomous or non-cell autonomous programs operate to regulate dendritic scaling vs. proportional growth and branching. With respect to non-cell autonomous regulation of dendritic scaling, studies have demonstrated that scaling growth of dendrites in *Drosophila* md neurons requires the microRNA *bantam*, which acts in epithelial cells to diminish Akt kinase activity in adjacent neurons and thereby regulate dendritic morphogenesis (Parrish *et al*., 2009). In terms of cell autonomous mechanisms for dendritic scaling, studies have demonstrated a role for the co-chaperone of HSP90, *CHORD*, which interacts with the TORC2 component, *Rictor*, to mediate dendritic scaling (Shimono *et al*., 2014). Our study reveals that *bdwf* LOF causes a proportional reduction in both dendritic length and branching such that the branch density remains unaffected, indicative of a neural requirement in promoting proportional dendritic growth and branching across multiple md neuron subtypes.

We identified at least two pathways whereby Bdwf functions to regulate dendrite morphogenesis, including transcriptional and translational regulation. With respect to transcriptional regulation, we characterized a regulatory relationship between the homeodomain TF Cut and Bdwf. Cut exhibits differential expression in morphologically diverse md neuron subtypes and is required to promote dendritic architecture in these neurons. The TFs Vestigial and Scalloped have been shown to limit Cut expression in the CII neurons (Corty, Tam and Grueber, 2016), whereas the TF Lola promotes its expression in CIV neurons (Ferreira *et al*., 2014). While the expression of Bdwf and Cut are independent of each other, our data indicate that these two proteins interact via a reciprocal feedback loop, wherein Cut acts upstream to promote Bdwf expression, while overexpression of Bdwf has a negative feedback effect thereby restricting the expression of Cut. Moreover, we demonstrate that Bdwf functions as a downstream effector of Cut-mediated dendritic morphogenesis thereby adding to the complex molecular and cellular processes via which Cut exerts control over cell-type specific dendritic development (Nanda *et al*., 2017; Jinushi-Nakao *et al*., 2007; Sulkowski *et al*., 2011; Iyer *et al*., 2012; Nagel *et al*., 2012; S.C. Iyer *et al*., 2013; Ferreira *et al*., 2014; Das *et al*., 2017; Clark *et al*., 2018; Bhattacharjee *et al*., 2022). Interestingly, the observation that Bdwf overexpression can negatively feedback to suppress Cut levels may provide some insight into the findings that both LOF and GOF for Cut produce similar dendritic hypotrophy phenotypes in CIV md neurons. Intriguingly, another BED domain bearing TF, Dref, has been shown to have an antagonistic regulatory relationship with Cut with respect to controlling PCNA gene expression (Seto *et al*., 2006).

With respect to translational roles for Bdwf, co-immunoprecipitation analysis, combined with mass spectrometry data, indicate that Bdwf interacts with numerous ribosomal proteins of both the large and small subunits. Mutations in ribosomal protein genes have previously been associated with growth defects in animals, leading to a “minute” phenotype (Marygold *et al*., 2007). Loss-of-function of ribosomal proteins generally leads to a defect in cellular growth, and knockdown of *RpL35A* has also been shown to cause cell growth and viability phenotypes when assayed in Kc_167_ and S2R^+^ cells (Boutros *et al*., 2004). In cultured mice hippocampal neurons, loss of ribosomal proteins, S6, S14, and L4 is associated with reduced protein synthesis that leads to a more simplified dendritic arbor (Slomnicki *et al*., 2016), while loss of *RpL22* led to a disruption in dendritic morphology in CIV md neurons (Lin *et al*., 2015). Mutations in *RpS24* are associated with Diamond-Blackfan Anaemia and in *Drosophila* has been shown to cause developmental delays which were rescued by expressing synaptic vesicle proteins in serotonergic neurons (Gazda *et al*., 2006; Deliu *et al*., 2022). The observation that *bdwf* mutation leads to a proportional reduction in dendritic growth and branching may be explained, at least in part, by the interaction of Bdwf with components of the ribosome, as knockdown of these genes leads to severe dendritic hypotrophy. Moreover, proper functioning of the nervous system requires neurons to respond to stimuli within minutes. Given the complexity of dendritic arbors, this is achieved through local protein translation at distal dendrites and axons (Holt, Martin and Schuman, 2019). The presence of ribosomes in dendrites and axons has long been established (Holt, Martin and Schuman, 2019; Singh *et al*., 2019; Lottes and Cox, 2020). Studies have also demonstrated that activity dependent local protein translation leads to remodeling of the actin cytoskeleton which in turn modulates dendritic spine morphology (Nakahata and Yasuda, 2018). Our data shows that loss of *bdwf* disrupts ribosome trafficking along the dendrites and in CIV neurons we demonstrate that *bdwf* LOF leads to reduced translation. With respect to the cytoskeleton, Cut has previously been shown to regulate dendritic architecture via downstream effectors that direct F-actin and MT dendritic cytoskeletal organization and stability (Jinushi-Nakao *et al*., 2007; Das *et al*., 2017; Bhattacharjee *et al*., 2022). Similarly, genetic depletion of *bdwf* or genes encoding ribosomal proteins destabilizes MT assembly as well as reduces F-actin rich dendritic branching, suggesting that Bdwf acts in concert with ribosomal proteins to promote dendritic growth by modulating both F-actin and MT cytoskeletal architecture.

We propose a model of nucleo-cytoplasmic Bdwf activity whereby the constellation of functional roles in ribosomal trafficking, protein translation, and cytoskeletal architecture, together with the TF regulatory relationship Bdwf shares with Cut, provides mechanistic insights for how this gene directs cell-type specific dendritic arborization (Figure 9). While both Cut and Bdwf are demonstrated to regulate the F-actin and MT cytoskeleton, whether this occurs as a direct mechanism of nuclear Bdwf awaits future investigation.

**Figure 9:**
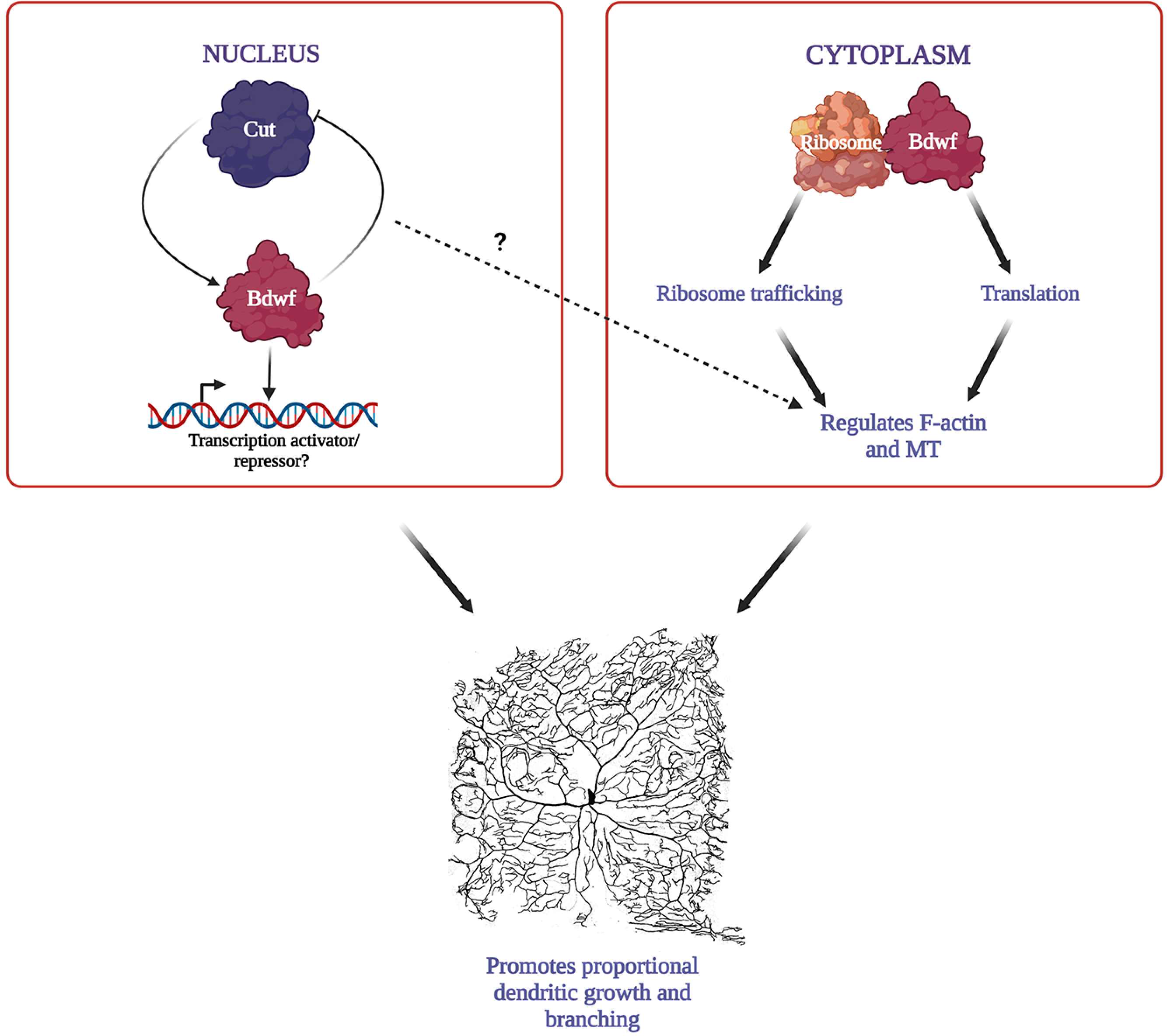
Model for Bdwf function in dendrite morphogenesis. Cut and Bdwf have a reciprocal regulatory relationship whereby Cut positively regulates Bdwf expression, and overexpression of Bdwf negatively regulates Cut expression. Bdwf has putative DNA binding activity. Bdwf interacts with ribosomal proteins to form ribonucleoprotein complex, which potentially impacts ribosome trafficking and protein translation. Bdwf modulates cytoskeletal structure by promoting MT and F-actin-dependent dendritic growth and branching in Cut-positive neurons. While both Cut and Bdwf regulate dendritic cytoskeletal components, whether this occurs as a direct mechanism of nuclear Bdwf is, as yet, unknown.

## Materials and Methods

### Drosophila strains

*Drosophila* stocks were raised on standard cornmeal-molasses-agar media at 25°C. Fly strains used in these studies were either generated (*UAS-bdwf*, *UAS-bdwf-BED-HA*, *UAS-bdwf-ΔBED-myc*) or obtained from Bloomington or Vienna *Drosophila* stock centers (Dietzl *et al*., 2007). *bdwf^d05488^* line was obtained from the Exelixis collection at Harvard Medical School (Thibault *et al*., 2004). The following gene-specific *UAS-RNAi* (*IR*) lines were tested: *bdwf-IR* (JF02831; GD10700); *cut-IR* (JF03304); *RpS24-IR* (v104676); *RpL4-IR* (v101346); *RpL22-IR* (v104506); *RpL31-IR* (v104467). *GAL4* strains for md neuron subtypes included: *GAL4^221^, UAS-mCD8∷GFP* (*CI-GAL4*); *ppkGAL4, ppkGAL80*, *UAS-mCD8∷GFP* (*CIII-GAL4*); *GAL4^477^,UAS-mCD8∷GFP,/CyO,tubP-GAL80* (*CIV-GAL4*); *GAL4^ppk1.9^,UAS-mCD8∷GFP* (*CIV-GAL4*) *GAL4^21-7^,UAS-mCD8∷GFP* (*pan-md-GAL4*). Other strains included: *UAS-cut*; *UAS-mCD8∷RFP*; *w, ct^c145^, FRT^19A^/y^+^, ct^+^, Y* (Grueber *et al*., 2003); *y,w,tubP-GAL80,hsFLP,FRT^19A^; GAL4^109(2)80^,UAS-mCD8∷GFP; UAS-RpL10Ab-GFP*; *UAS-dMetRS^L624G^-3xmyc* (Erdmann *et al*., 2015); *UAS-Luciferase-IR; UAS-GMA∷GFP;GAL4^477^, UAS-mCherry∷Jupiter* (Das *et al*., 2017); *GAL4^Act5c^; UAS-CD4-tdTomato*. *Oregon-R* was used as a wild-type strain. UP-TORR (https://www.flyrnai.org/up-torr/) was used to computationally assess predicted off-target effects for gene-specific RNAi constructs (Hu *et al*., 2013) and all IR lines used have no predicted off-target effects.

### Transgenic strain generation

For this study, we generated full-length Bdwf overexpression transgene (*UAS-bdwf-FLAG-HA*) as well as structure-function variants (*UAS-bdwf-BED-HA* and *UAS-bdwf-ΔBED-myc*). To generate the full length overexpression transgene, a full-length cDNA for the sole *bdwf* mRNA isoform (BDGP clone LD44187) was subcloned into *pUAST* with a C-terminal insertion of FLAG and HA epitope tags. For structure-function variants, we custom synthesized versions of *bdwf* (Genscript, Piscataway, NJ, USA) that express only the BED domain (amino acid 1-47) or that lack the BED domain (ΔBED; removal of amino acids 6-47). These custom gene synthesis products were then Gateway® subcloned into the pTWH plasmid for the BED only domain or into the pTWM plasmid for the ΔBED variant. pTWH is a *pUAST* vector bearing a C-terminal HA epitope tag, while pTWM is a *pUAST* vector bearing a C-terminal myc epitope tag. All three *bdwf* transgenes were generated by ΦC31-mediated integration with targeting to 2R (51C1) (BestGene, Chino Hills, CA, USA). All constructs were sequenced verified for accuracy.

### Cell Isolation

The isolation and purification of CI da neurons was performed as previously described (Iyer *et al*., 2009). Briefly, 40-50 age-matched third instar larvae having the genotypes of either +/+; *GAL4^221^,UAS-mCD8∷GFP* or *UAS-Cut/+; GAL4^221^,UAS-mCD8∷GFP* were collected and washed several times in ddH20. The larvae were then rinsed in RNAse away, ddH20 and finally dissected. The tissue was then dissociated using a combination of enzymatic and mechanical perturbations to yield single cell suspensions which were filtered using a 30μm membrane. The filtrate is then incubated with superparamagnetic beads (Dynabeads MyOne Streptavidin T1, Invitrogen) coupled with biotinylated mouse anti-CD8a antibody (eBioscience) for 60 minutes. Finally, the CI neurons attached to the magnetic beads were then separated using a powerful magnetic field. The isolated neurons were washed at least five times to remove any potential nonspecific cells and the quality and purity of isolated neurons was assessed under a stereo-fluorescent microscope equipped with phase contrast for examining the number of fluorescent/nonfluorescent cells. Only if the isolated cells were free of cellular debris and non-specific (i.e. non-fluorescing) contaminants where they retained. The purified CI neurons were lysed in RNA lysis buffer and stored until needed for qRT-PCR analyses.

### qRT–PCR analysis

qRT–PCR analysis of control and Cut overexpressing CI da neurons was performed in triplicates, and repeated twice independently. The expression levels of *cut* and *bdwf* were assessed by qRT–PCR. Values obtained from these analyses were normalized to the endogenous control (*RpL32*), and the levels relative to those observed in flow through fraction were calculated using the ΔΔCτ method (Livak and Schmittgen, 2001). Pre-validated Qiagen QuantiTect Primer Assays (Qiagen, Germantown, MD, USA) were used for *cut* (QT00501389), *bdwf* (QT00976549) and *RpL32* (QT00985677).

### Phenotypic screening, live image confocal microscopy, and morphometric quantitation

Live imaging was performed as previously described (Iyer *et al*., 2013). Briefly, *GAL4^477^,UAS-mCD8∷GFP*, *GAL4^ppk^*, or *GAL4^ppk^,GAL80^ppk^,UAS-mCD8∷GFP*, or *GAL4^221^,UAS-mCD8∷GFP* female virgins were aged for 2-3 days and crossed to *UAS-bdwf-IR* or *UAS-bdwf-FLAG-HA* transgenic males, and reared at 29°C. *Oregon R* flies were used as controls. For Bdwf-Ct interaction in CI neurons, *UAS-ct; GAL4^221^,UAS-mCD8∷GFP* female virgin flies were crossed to either *UAS-ct*, or *UAS-bdwf-IR*, or *UAS-mCD8∷RFP*, and *Oregon R* was used as controls. To analyze Bdwf-Ct interaction in CIV neurons, *GAL4^477^,UAS-mCD8∷GFP; ct-IR* virgin female flies were crossed to either *UAS-CD4-tdTomato* (B35837) or *UAS-bdwf-IR*. *GAL4^477^,UAS-mCD8∷GFP* virgin female flies that were outcrossed to *UAS-CD4-tdTomato* were used as controls. Images were acquired from at least 6-10 wandering third instar larvae. For live image analyses, larvae were placed on a microscope slide, immersed in 1:5 (v/v) diethyl ether to halocarbon oil and covered with a 22 × 50 mm glass coverslip. Neurons expressing GFP were visualized on either a Nikon C1 Plus confocal microscope or a Zeiss LSM 780. For Nikon C1 plus microscope, images were collected as z-stacks using a 20X oil immersion lens at a step-size of 2.5 μm and 1024 × 1024 resolution. For LSM 780, images were collected as z-stacks using a 20X dry lens at a step-size of 2 μm and 1024×1024 resolution. The images were then converted to maximum intensity projections and were processed as previously described using ImageJ (Das et al., 2017; Iyer et al., 2013; Sulkowski et al., 2011). Quantitative morphometric data such as total dendritic length and number of branches were extracted and compiled using a custom Python script and the data output was imported into Microsoft Excel. Morphometric data was analyzed in Microsoft Excel and statistical tests were performed and graphs were plotted using GraphPad Prism 9.

For the Bdwf protein interactor cytoskeletal screen, *UAS-GMA;GAL4^477^,UASmCherry∷Jupiter* (Das et al., 2017) virgin flies were outcrossed to either *UAS-bdwf-IR, UAS-ct-IR* (B33967), *UAS-RpS24-IR* (v104676), *UAS-RpL4-IR* (v101346), *UAS-RpL22-IR* (v104506), *UAS-RpL31-IR* (v104467) transgenic male flies or *Oregon R* males. Images were acquired from wandering third instar larvae on a Zeiss LSM780 confocal microscope using a 20X dry lens objective at a step-size of 1.5μm and 1024×1024 resolution. To analyze ribosome localization, *UAS-RpL10Ab∷GFP; GAL4^221^,mCD8∷RFP*, or *UAS-RPL10Ab∷GFP; GAL4^ppk1.9^,UAS-mCD8∷RFP* virgin female flies were crossed to *Oregon R* (control) or *UAS-bdwf-IR*. For ribosome localization, images were acquired at 63x and a step size of 1 μm and 1024 × 1024 resolution.

MARCM analyses were performed as previously described (Grueber *et al*., 2003; Sulkowski *et al*., 2011; Das *et al*., 2017). Briefly, *w, ct^c145^, FRT^19A^/y^+^, ct^+^, Y* males were outcrossed to *y,w,tubP-GAL80,hsFLP,FRT^19A^; GAL4^109(2)80^,UAS-mCD8∷GFP* virgin females. Late third instar larvae (96-120 hr after egg lay) were then examined for the presence of GFP labeled *cut* mutant clones, followed by dissection, fixation, and staining as described below for immunohistochemistry.

### Multichannel reconstructions

Multichannel cytoskeletal reconstructions and quantitative analysis was performed using a previously described method (Nanda *et al*., 2021). Ribosome signal was quantified following a similar procedure. Two-channel neuronal image stacks with membrane and ribosomal expression were processed to create multi-signal reconstructions (Nanda *et al*., 2018). The local expression for dendritic compartment was quantified as follows:

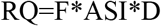

Here, RQ is the local ribosomal quantity, F is the fraction of the dendritic compartment volume occupied by the ribosomal signal, ASI is the average signal intensity, and D is local dendrite thickness, which is approximated by measuring the local dendritic diameter from the reconstructions. The average ribosomal expression per micrometer was then calculated for each individual neuron by dividing total ribosomal quantify by total length. Average expression specifically at the branch points were also calculated for each neuron. Both overall and branch level average expression levels for all neurons were normalized, and were expressed as a fraction of the maximal average expression found in all neurons. All reconstruction data have been uploaded to NeuroMorpho.Org (doi: 10.1038/sdata.2018.6) and will be released in the Cox archive upon publication.

### Generation of anti-Bdwf antibodies

The Bdwf protein sequence was obtained from flybase.org (CG3995-PA) and analyzed by a Hopp and Woods Antigenicity Plot to predict high surface probability of putative Bdwf antigens that lie outside of conserved Zinc-BED domain and show protein sequence specificity unique to Bdwf protein. Based upon these analyses, a distinct 15 amino acid peptide sequence (EFDVDEVEDVVPEED) in the middle of exon 2 was conjugated to KLH to improve antigenicity of the peptide sequence followed by polyclonal antibody production in rabbits (Genscript, Inc. Piscataway, NJ, USA). Anti-Bdwf antibody was affinity purified against the antigenic peptide and specificity was confirmed by immunohistochemistry analyses.

### Immunohistochemistry

Dissection, staining, and mounting of wandering third instar larvae was performed as previously described (Sulkowski *et al*., 2011). For immunohistochemistry (IHC), larvae were dissected in PBS, pinned on sylgard plates and fixed in 4% paraformaldehyde for 20 minutes. Next, larvae were washed five times in 1X PBT (PBS + 0.3% Triton X-100). Cuticle filets were blocked in 5% normal donkey serum (Jackson Laboratories, West Grove, PA) for at least 30 minutes at room temperature followed by incubation with respective primary antibodies. Antibody dilutions used were as follows: rabbit anti-Bdwf (1:200), mouse anti-CD8 (1:100) (Invitrogen), rabbit RpS6-(1:100) (Cell Signaling Technology), and mouse anti-Cut (2B10; 1:50) (Developmental Studies Hybridoma Bank). Donkey anti-rabbit and donkey anti-mouse secondary antibodies were used at 1:200 (Jackson Immunoresearch). Filets were incubated in glycerol for 5 minutes, followed by being directly mounted onto coverslips with either a drop of glycerol, or aqueous fluoromount. The slides were then imaged on a Nikon C1 Plus confocal microscope or Zeiss LSM 780.

### FUNCAT analysis

FUNCAT analysis was performed as previously described (Erdmann *et al*., 2015, 2017; Marter *et al*., 2019). Briefly, flies expressing *UAS-dMetRS^L624G^-3xmyc* under the control of *GAL4^ppk1.9^* were crossed to either *UAS-Luc-IR* (control) or *UAS-bdwf-IR* and grown in blue-food containing 4mM azidonorleucine (ANL) (Torcis, Cat No 6585) and incubated at 29°C. Wandering third instar larvae with visible blue dye in the guts were selected and dissected and fixed at 4% paraformaldehyde for 20 minutes. Larval fillets were washed with 1X PBT three times followed by three washes with 1X PBS (pH 7.4) for 15 minutes each. Following washes, the fillets were treated with 1X PBS (pH 7.4) solution containing triazole ligand (200 mM TBTA, Sigma-Aldrich 1:1000 dilution), TAMRA-alkyne dye (Invitrogen, 1 mM, 1:5000 dilution), tris-(2-carboxyethyl) phosphine (400 mM, Sigma-Aldrich 1:1000 dilution), and CuSO_4_ (200 mM, Sigma Aldrich, 1:1000) and incubated overnight at 4 °C with gentle agitation. The fillets were then washed with 1X PBST (1X PBS pH 7.4 + 1% v/v Tween-20) the following day for 15 minutes each and then incubated with Alexa-Fluor goat anti-horseradish peroxidase (HRP) 488 (1:200) overnight at 4 °C. Following immunostaining the fillets were washed in 1X PBT twice and once in 1X PBS (pH 7.4) and mounted in aqueous fluoromount and imaged on Zeiss LSM 780 at 63X objective.

### Mass Spectrometry Analysis

Larval homogenates were prepared as described in Iyer *et al*. (2009). Age matched third-instar larvae expressing *UAS-bdwf-FLAG-HA* driven by the ubiquitous *GAL4^Act5C^* driver were homogenized. Cell pellets were lysed in non-denaturing lysis buffer with protease inhibitors (50 mM Tris/HCl pH 7.5, 1 mM EGTA, 1 mM EDTA, 0.5 or 1% (v/v) NP-40, 1 mM sodium orthovanadate, 50 mM NaF, 5 mM sodium pyrophosphate, 0.27 M sucrose, 10 mM sodium 2-glycerophosphate, 0.2 mM phenylmethylsulphonyl fluoride, 1 mM benzamidine, plus 100 mM iodoacetamide added fresh prior to lysis, and pepstatin/aprotinin to inhibit proteases). Cell lysates were clarified by centrifugation at 14,000 × g for 15 min at 4°C. The supernatant was collected and protein concentration determined. Cell lysates were incubated with anti-HA tag antibody overnight. Bead only or IgG only controls were performed in parallel to control for specificity (Table S1). To capture protein of interest, 1 mg of cell extract protein was used and incubated for 3 h to overnight at 4°C with the affinity resin. The beads are then washed three times with 1 ml of lysis buffer containing 500 mM NaCl and once with 0.5 ml of 10 mM Tris/HCl pH 8.0.

The soluble protein samples from individual samples were dried with SpeedVac, reconstituted in 8 M urea, reduced by 10 mM DTT for 30 min, alkylated by 50 mM iodoacetamide for 30 min, and digested by trypsin at 37°C overnight. Tryptic peptides were further purified by Zip-Tip (Millipore) and analyzed by LC-MS/MS using a linear ion-trap mass spectrometer (LTQ, Orbitrap). Subsequent to sample injection, the column was washed for 5 min with mobile phase A (0.4% acetic acid) and peptides eluted using a linear gradient of 0% mobile phase B (0.4% acetic acid, 80% acetonitrile) to 50% mobile phase B in 30 min at 250 nl/min, then to 100% mobile phase B for an additional 5 min. The LTQ mass spectrometer was operated in a data-dependent mode in which each full MS scan was followed by five MS/MS scans where the five most abundant molecular ions were dynamically selected for collision-induced dissociation using a normalized collision energy of 35%. Tandem mass spectra were collected by Xcalibur 2.0.2 and searched against the NCBI *Drosophila* protein database using SEQUEST (Bioworks 3.3.1 software from ThermoFisher) using tryptic cleavage constraints. Mass tolerance for precursor ions was 5 ppm and mass tolerance for fragment ions was 0.25 Da. SEQUEST filter criteria were: Xcorr vs. charge 1.9, 2.2, 3.5 for 1+, 2+, 3+ ions; maximum probability of randomized identification of peptide <0.01. Protein identifications and number of identifying spectra (peptide hits) for each sample were exported using >99% confidence limit for protein identification with peptides from a given protein identified at least in two independent samples (Table S1). Functional enrichment analysis was performed using DAVID (Dennis *et al*., 2003; Huang, Sherman and Lempicki, 2008) to identify statistically overrepresented functional gene classes. For these analyses the protein-interactors of Bdwf were used as input and all genes in the genome were selected as background. DAVID uses the Fisher’s Exact statistics to retain significant results calculated after redundancy in original gene lists are removed (Table S1). The EASE Score P-value ranges from 0 to 1 where a value of 0 represents perfect enrichment. P-values equal or smaller than 0.05 were considered strongly enriched in the annotation categories.

## Acknowledgements

This work was supported by NIH R01 NS086082 (DNC/GAA) and NIH R01 NS039600 (GAA). We thank the TRiP at Harvard Medical School (NIH/NIGMS R01-GM084947) for providing transgenic RNAi fly stocks and/or plasmid vectors used in this study. We thank Dr. Daniela C. Dieterich (University Magdeburg, Germany) for providing the *UAS-dMetRS^L624G^-3xmyc* transgenic strain and thank Dr. Binghe Wang (Georgia State University, Atlanta, GA) for consulting on the FUNCAT analyses. We thank the Georgia State University Imaging Core Facility for personnel training and instrument support. We acknowledge Emily Wilde, Luis Sullivan, Shaurya Prakash, Yukting Lau, Vihitha Thota, and Farheen Shaikh from the Cox Lab for assistance with imaging, genetics, and neuronal reconstructions reported in this study.

## Author Contributions

SB, EPRI, and DNC designed the study with computational inputs from SN and GAA. SB, EPRI, SCI, SN, MR performed the formal analyses and investigations. SB, EPRI, DNC wrote the original draft. SB, SN, GAA, and DNC performed review and editing. GAA and DNC were responsible for funding acquisition and supervision.

---

***Figure S1: Rescue of bdwf mutant dendritic defects and validation of bdwf-IR and Bdwf antibody***. (A-E) Compared to wild-type control CIV neurons (A), *bdwf ^d0588^* homozygous mutant CIV neurons results display a strong and penetrant dendritic hypotrophy (B), which is fully rescued by CIV expression of *UAS-bdwf* (C, D, E). Genotypes: (A-E) *Gal4^477^, UAS-mCD8∷GFP/+; +/+* or *bdwf ^d0588^/ bdwf ^d0588^* or *UAS-Bdwf-FLAG-HA/ +*. Scale bar, 50 μm, n=8 neurons per genotype. (F-H) *bdwf-IR* knockdown in CIV neurons leads to a significant reduction in Bdwf protein expression levels. Scale bar, 10 μm. n= 16 (control) and n=9 (*bdwf-IR)*. Error bars represent +/− S.E.M. ***p≤0.01 (Student’s t-test).

***Figure S2: Cut is not required for Bdwf expression and Bdwf can partially rescue cut RNAi-mediated defects in CIV dendrite morphogenesis**.* (A-C) Representative images of *ct* MARCM clones of CIV md neurons stained if CD8 (green) and Bdwf (magenta) showing normal Bdwf expression in *ct* mutants. Scale bar: 1 μm. (D-F) Bdwf acts downstream of Ct to promote dendritic growth in CIV md neurons. Compared to control (D), *ct-IR* (E, G, H) shows significant reduction in both the total dendritic length and the total dendritic branches which is partially rescued by the simultaneous co-expression of *UAS-bdwf-FLAG-HA* and *ct-IR* (F, G, H) in CIV ddaC neurons. Scale bar, 100 μm. (G, H) Quantification of TDL and TDB for CIV da neurons. Error bars represent +/− SEM (One-way ANOVA with Sidak’s multiple comparison test). n=9 for WT, n=8 for *ct-IR*, n=8 for *bdwf-OE+ct-IR*. Genotypes: *GAL4^477^,UAS-mCD8∷GFP/+;UAS-CD4-tdTOM/+* (D, G, H), *GAL4^477^,UAS-mCD8∷GFP/+;UAS-ct-IR/ UAS-CD4-tdTOM* (E, G, H), *GAL4^477^,UAS-mCD8∷GFP/+;UAS-ct-IR/ UAS-bdwf-FLAG-HA* (F, G, H).

***Table S1**: **Mass spectrometry and functional annotation of Bdwf protein interactors***. Summary of Bdwf protein-protein interactors isolated by immunoprecipitation and characterized by mass spectrometry, including affinity bead only and IgG only controls. Summary of DAVID functional annotation analyses of gene ontologies enriched in the Bdwf protein-protein interaction dataset.

