## Supplementary figures and images for "The Zinc-BED transcription factor Bedwarfed promotes proportional dendritic growth and branching through transcriptional and translational regulation in *Drosophila*"

### Figure S1

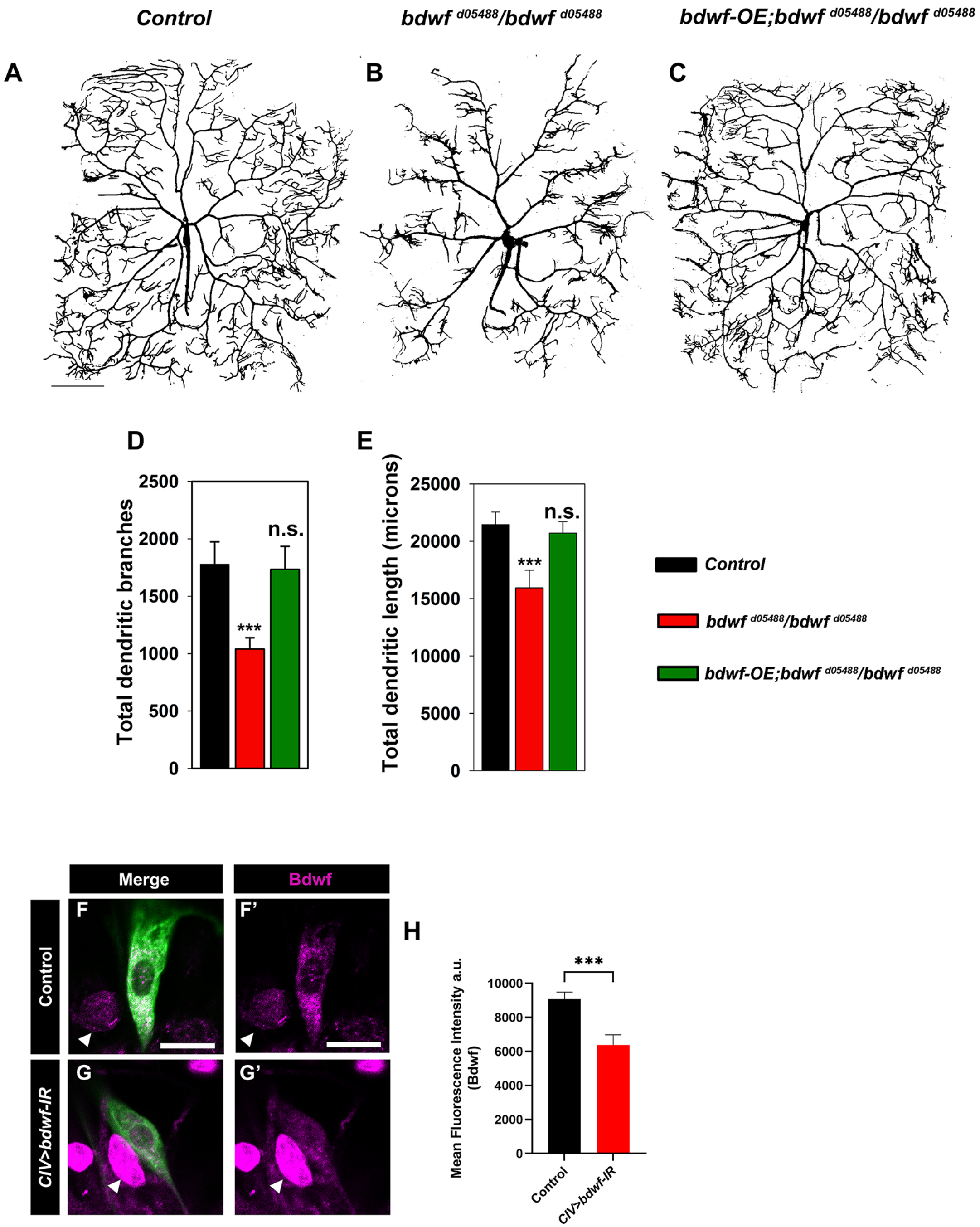

### Figure S2

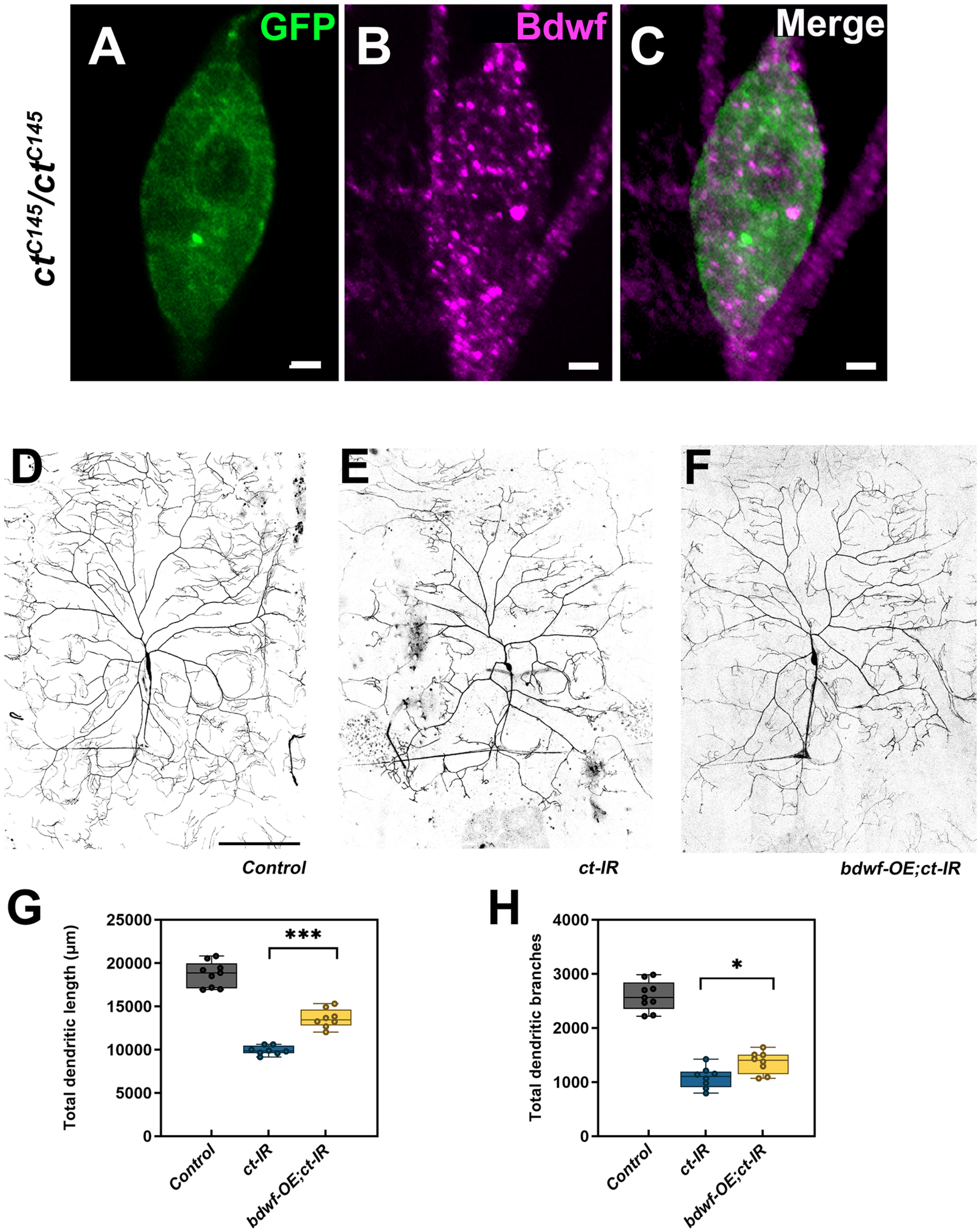
